# tRNA 2’-O-methylation modulates small RNA silencing and life span in *Drosophila*

**DOI:** 10.1101/699934

**Authors:** Margarita T. Angelova, Dilyana G. Dimitrova, Bruno Da Silva, Virginie Marchand, Catherine Goyenvalle, Cyrinne Achour, Caroline Jacquier, Valérie Bourguignon-Igel, Salman Shehzada, Tina Lence, Vincent Guerineau, Jean-Yves Roignant, Christophe Antoniewski, Laure Teysset, Damien Bregeon, Matthias R. Schaefer, Yuri Motorin, Clément Carré

## Abstract

2’-O-methylation (Nm) represents one of the most common RNA modifications. Nm affects RNA structure and function with crucial roles in various RNA-mediated processes ranging from RNA silencing, translation, self *versus* non-self recognition to viral defense mechanisms. Here, we identify two novel Nm methyltransferases (Nm-MTases) in *Drosophila melanogaster* (CG7009 and CG5220) as functional orthologs of yeast TRM7 and human FTSJ1, respectively. Genetic knockout studies together with MALDI-TOF mass spectrometry and RiboMethSeq mapping revealed that CG7009 is responsible for methylating the wobble position in tRNA^Phe^, tRNA^Trp^ and tRNA^Leu^, while subsequently, CG5220 methylates position C32 in the same tRNAs and targets also additional tRNAs. *CG7009* or *CG5220* mutant animals were viable and fertile but exhibited various phenotypes such as life span reduction, small RNA pathways dysfunction and increased sensitivity to RNA virus infections. Our results provide the first detailed characterization of two TRM7 family members in *Drosophila* and uncover a molecular link between enzymes catalysing Nm at specific tRNAs and small RNA-induced gene silencing pathways.

## Introduction

The existence of RNA modifications has been known for over 50 years and many of the pioneering studies addressed the function of RNA modifications in abundantly expressed RNAs such as transfer RNAs (tRNAs) and ribosomal RNAs (rRNAs). tRNAs are the most heavily modified RNAs (1). tRNAs are modified post-transcriptionally and the biosynthesis of modified nucleosides often requires different modification enzymes acting sequentially at distinct steps of tRNA maturation (2, 3). The complex mechanisms underlying the stepwise modification of tRNAs were largely deciphered in the yeast *S. cerevisiae*, as well as in studies conducted in prokaryotes and *Archaea*. More recently, some of these seminal findings in single-cell organisms were revisited using multi-cellular models including studies that aim at understanding how mutations in tRNA modification enzymes affect organismal development and disease etiology.

2’-O-methylation (Nm) is a common RNA modification. The addition of a methyl group to the 2’ hydroxyl of the ribose moiety of a nucleoside creates Nm (reviewed in (4, 5)). Nm can occur at any nucleotide explaining the abundant nature of this modification. Nm residues are found at multiple and often highly conserved positions in tRNAs, rRNAs, and small nuclear RNAs (snRNAs) (6–8). In eukaryotes, RNA modification reactions resulting in Nm on rRNAs and snRNAs are frequently catalyzed by evolutionarily conserved C/D-box small RNAs (SNORDs) involving guide ribonucleoprotein particles (RNPs) which contain the Nm-methylase fibrillarin. Small nucleolar RNPs (snoRNPs) mediate the deposition of Nm at rRNAs while small Cajal bodies RNPs (scaRNPs) direct Nm-modification to snRNAs. In contrast, most of the Nm deposition occurring in eukaryotic tRNAs are mediated by stand-alone proteins without the need for guidance by small RNAs. However, recently it was reported that one snoRNA and one scaRNA can guide Nm deposition to tRNA^Met^ in mammalian cells (9). Importantly, Nm deposition was also reported at 3’-terminal nucleotides in small non-coding RNAs (sncRNAs) such as microRNAs (miRNAs) and small-interfering RNAs (siRNAs) in plants (10–12), in Argonaute-2 (Ago2) loaded siRNAs and miRNAs in flies and in PIWI-interacting RNAs (piRNAs) in animals (13–15). More recently Nm was also reported to be present in messenger RNA (mRNA) (16, 17).

Nm can affect RNAs in multiple ways as it increases hydrophobicity, protects RNAs from nuclease degradation (9, 15, 18), stabilizes helical structures or modulates interactions with proteins or other RNAs (19–30).

The majority of all tRNA modifications are deposited at the wobble position N_34_ in the anticodon loop (ACL), and at the anticodon-adjacent position N_37_. Amongst the different tRNA isoacceptors, these two positions contain highly conserved modifications, which is suggestive of their physiological importance. Accordingly, it was shown that ACL modifications prevented frameshifting during translation (31, 32) and are thus necessary for the correct decoding of genetic information (33).

The methyltransferase complex catalysing Nm formation in the ACL of mammalian and yeast tRNAs comprises the Nm-methyltransferases (Nm-MTases) FTSJ1 or TRM7, respectively. These enzymes associate with specific proteins: THADA/TRM732 for Nm_32_ and WDR6/TRM734 for Nm_34_ formation (2, 34, 35). Cm_32_ and, more importantly Nm_34_, are required for efficient formation of a third modification, wybutosine (yW) at m^1^G_37_ in tRNA^Phe (^36–38^)^. The same circuitry is conserved in the phylogenetically distant yeast *Schizosaccharomyces pombe* (35) while the formation of peroxywybutosine (o2yW) at position 37 is also affected in humans lacking FTSJ1 (35, 39).

Several studies have uncovered crucial roles for FTSJ1/TRM7 in normal and pathological conditions (reviewed in (4, 5)). While in *S. cerevisiae*, lack of TRM7 affected growth, *FTSJ1* mutant mice showed impairment in their learning capacity as well as significantly reduced pain sensing (hypoalgesia) and altered gene expression profiles (40). Similarly, in humans several mutations in *FTSJ1* were shown to be causative of a neurodevelopmental disorder known as Non-Syndromic X-linked Intellectual Disability (NSXLID) (39, 41, 42). Importantly, expression of human FTSJ1 in yeast suppressed the severe growth defects observed in *trm7Δ* mutants, demonstrating that the TRM7 enzyme family and their RNA targets are highly conserved (35).

While the molecular function of yeast and human Nm-MTases acting on specific tRNAs has been established, the molecular mechanisms causing the complexity of observed mutant phenotypes have not fully been elucidated. Importantly, a tractable multicellular model system that would allow studying Nm-MTase function systematically and thereby bridge the growth phenotypes observed in *trm7* deficient yeast with the complex phenotypes observed in FTSJ1-mutant man has been lacking.

In this report, we show that, in contrast to yeast and humans, *Drosophila melanogaster* has evolved two Nm-MTase genes, *CG5220* and *CG7009*, whose genes products specialized their activity to respectively methylate positions 32 and 34 in the ACL of specific tRNAs. We demonstrate that the catalytic specificity of these Nm-MTases is dependent on the position rather than the identity of the ACL nucleotides. Importantly, lack of these proteins reduced *Drosophila* life span and impaired various cellular pathways, which employ small RNAs to achieve post-transcriptional silencing. Hence, *CG5220* and *CG7009* mutant animals were more sensitive to RNA virus infections and showed dysfunctional control of transposable elements, suggesting a molecular link between Nm RNA modifications and small RNA gene silencing pathways in *Drosophila*.

## Results

### A genome-wide RNAi screen identifies CG7009 as regulator of siRNA-mediated silencing pathways

We previously developed and characterized a self-silencing genetic sensor (*automiG*) that combines the expression of GFP with miRNAs, miG-1 and miG-2, targeting GFP mRNA (Figure 1A and (43)). *AutomiG* self-silencing reports on the activity of canonical miRNA biogenesis factors such as Drosha and Dicer1 (Dcr1) and the function of RNA-induced silencing complex (RISC) factors such as Argonaute2 (Ago2) and Dicer2 (Dcr2). Impairing the function of miRNA biogenesis or Ago2 silencing activity thus causes the de-repression of *automiG* self-silencing resulting in the expression of GFP (43). To identify additional regulators of these two RNA silencing pathways, a genome-wide RNA interference screen was performed in *Drosophila* S2 cells expressing the *automiG* sensor. Using a double-stranded RNA (dsRNA) collection library (the DRSC 2.0) allowed the down-regulation of 94.3% of all annotated *Drosophila* genes. The screen identified known regulators of miRNA biogenesis such as Drosha and Pasha as well as siRNA pathway silencing key actors like Ago2 and Droj2 (Supplementary Figure S1A), demonstrating the validity of this approach. In addition, we identified 17 additional genes affecting a*utomiG* silencing, which had not been reported to act in siRNA and/ or miRNA pathways (Supplementary Figure S1A). Among those, CG7009 stood out as an uncharacterized gene with sequence identity to annotated Nm-MTases. RNAi-mediated inactivation of CG7009 in S2 cells expressing *automiG* resulted in increased GFP expression when compared to control constructs (Figure 1B). In addition, a dual luciferase assay reporting on siRNA pathway activity in S2 cells (44, 45) confirmed that Dcr2/Ago2-dependent silencing was affected in cells with down-regulated CG7009 expression (Supplementary Figure S1B).

**Figure 1.**
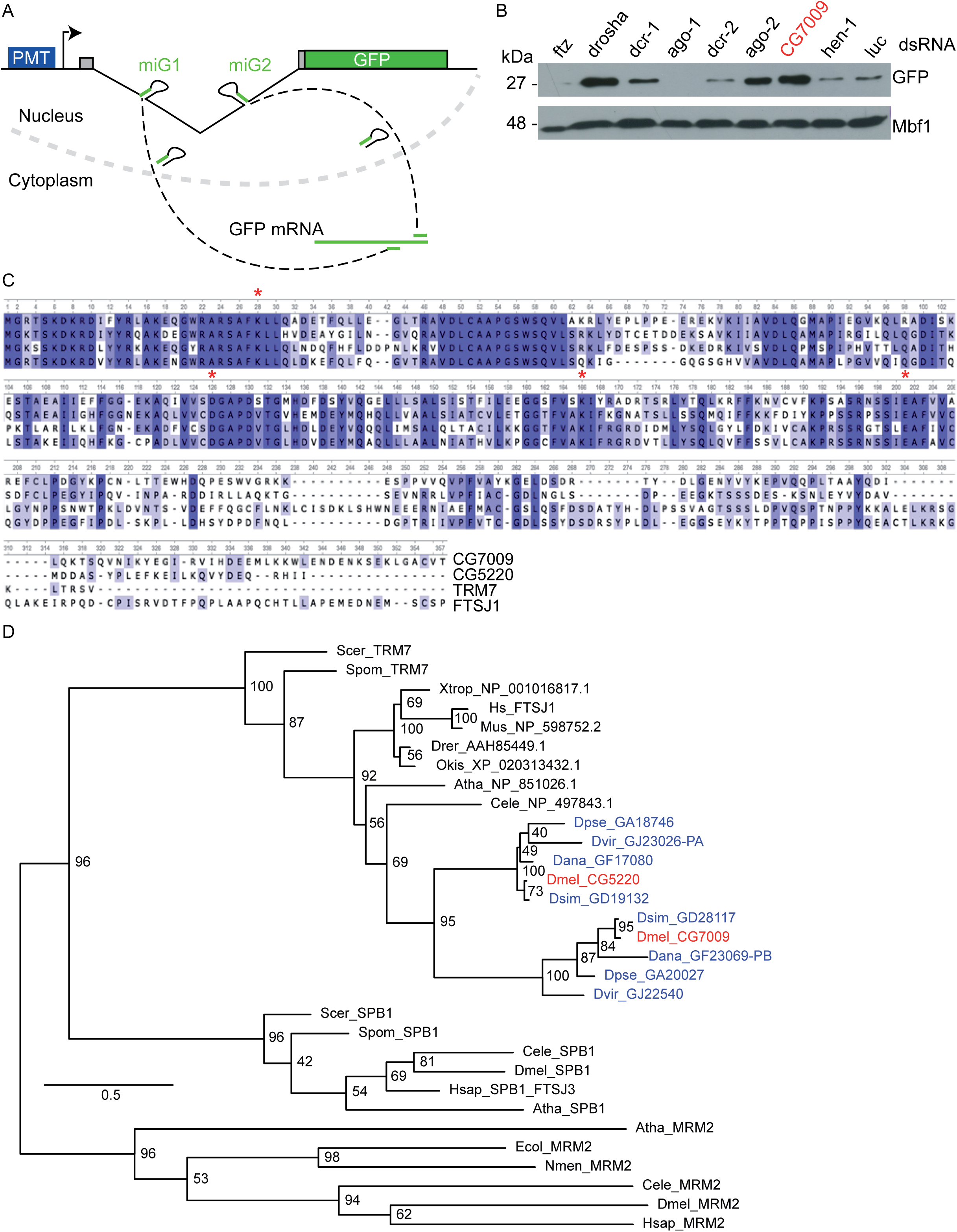
Identification of CG7009 as a conserved TRM7 family protein. **A**, The *automiG* sensor. *automiG* carries a copper inducible promoter (PMT) that drives the expression of two miRNAs (miG1 & miG2) and the GFP mRNA. Both miRNAs target the GFP mRNA with perfect complementarity. *AutomiG* repression is dependent on Ago2, Drosha, Pasha, Dicer-1 and Dicer-2 functions (43). **B**, CG7009 function affects *automiG* repression. Cells were soaked with the indicated double-stranded (dsRNA), followed by *automiG* induction. GFP expression was analyzed by western blotting. The Mbf1 protein was used as a loading control. *Ftz* (*Fushi tarazu*) and *luc* (luciferase) dsRNA served as negative KD control. KDa: Kilo Dalton. **C**, Multiple amino acid sequence alignment of CG7009, CG5220, TRM7, and FTSJ1. The conserved predicted catalytic tetrad K-D-K-E is marked with red asterix (*). **D**, *Drosophila* species evolved two TRM7 family proteins. Phylogenetic analysis of TRM7 and SBP1 MTases. The RrmJ SBP1 family of Nm-MTases that act on rRNA was used as an outgroup. Blue color blue indicates *Drosophila* species proteins. Red indicates *Drosophila melanogaster* proteins.

In order to obtain insights into the impact of CG7009 loss on gene expression control through Dcr2/Ago2-mediated post-transcriptional gene silencing, we performed a transcriptome analysis in *Drosophila* S2 cells upon KD of CG7009 expression. Surprisingly, the KD of *CG7009* led to the deregulation of only 110 genes (FDR <0.01). Strikingly, the most statistically significant de-regulated gene (40% decrease, log2FC −0.7, FDR-adjusted *p*-value 7.73e^−118^) was the *Ago2* mRNA (Supplementary Figure S1C) suggesting that CG7009 may act upstream in the siRNA pathway by regulating Ago2 mRNA levels.

This genetic screen thus identified CG7009, a potential Nm-MTase involved in Dcr2/Ago2-mediated post-transcriptional silencing.

### CG7009 encodes for a predicted Nm-MTase

Amino acid (aa) sequence analysis suggested that the protein encoded by CG7009 in *D. melanogaster* harbours a methyltransferase domain belonging to the conserved RlmE family and TRM7 subfamily of the class I-like SAM-binding methyltransferase superfamily (38). Sequence alignment of the putative CG7009 protein with the yeast Nm-MTase TRM7 showed 52 % aa sequence identity, including the conserved KDKE motif in the active site, and 66 % of aa coverage (Figure 1C). FTSJ1 is the human ortholog of TRM7 (39). CG7009 shares 51 % aa identity and 86 % aa coverage with FTSJ1 (Figure 1C). Surprisingly, further sequence alignment of CG7009 protein with proteomes of different *Drosophila* species uncovered an additional gene, CG5220, whose annotated protein in *Drosophila melanogaster* displays 63 % aa sequence identity with CG7009 (Figure 1 C, D). Like CG7009, CG5220 was an uncharacterized gene with an amino acid composition that clearly showed an Nm-MTase signature (Figure 1C). Importantly, it was previously reported that overexpression of CG5220 was able to rescue the growth phenotype observed in *trm7Δ* mutant yeast (35). Similarly to CG7009, CG5220 shows high similarity to TRM7 (48 % identity and 83 % coverage) as well as to FTSJ1 (58 % of identity and 82 % coverage, Figure 1C). These findings pointed to CG7009 and CG5220 as potential paralogs and conserved members of the TRM7 Nm-MTases family in *Drosophila*.

### Mutations in CG7009 or CG5220 are viable and fertile

To investigate the function of CG7009 and CG5220 during *Drosophila* development and to characterize the potential enzymatic activity of their gene products, we characterized existing mutations in *CG7009* gene and generated *CG5220* mutant flies. For *CG7009*, one transposon insertion line (*CG7009^e02001^*) and two genomic deletion lines (*Def3340* and *Def9487*) were obtained and confirmed at the molecular level (Supplementary Figure S2A-E). Both *CG7009^e02001^* homozygous mutants and trans-heteroallelic combinations with both deficiencies were incapable of transcribing *CG7009* properly. In addition, a transgenic rescue line (called BAC throughout the manuscript) containing the *CG7009* genomic locus was established through BAC transgenesis (46) resulting in an insertion of ∼20 kb genomic sequence in an ectopic genomic location (Supplementary Figure S2D, E). To address the function of CG5220, CRISPR/Cas9-mediated genome editing was used to create a *CG5220* mutant allele (*CG5220^K>A^*), which substituted a conserved lysine at position 28 in the predicted catalytic domain with alanine (Figure 1C and Supplementary Figure S2F). The same substitution was reported to abolish the catalytic function of both yeast TRM7 and human FTSJ1 (39). Flies homozygous for either *CG7009^e02001^* or *CG5220^K>A^* or trans-heterozygous *CG7009* mutants, as well as *CG7009^e02001^*, *CG5220^K>A^* double mutants survived until adulthood under standard conditions. We observed neither a major growth defect as reported for yeast (36, 38) nor significant developmental delays of flies homozygous for either *CG7009^e02001^* or *CG5220^K>A^* or trans-heterozygous *CG7009* mutants. However, double mutants flies that were homozygous for *CG7009^e02001^*,*CG5220^K>A^* showed a mild reduction of size and weight when compared to controls (Figure 2A).

**Figure. 2.**
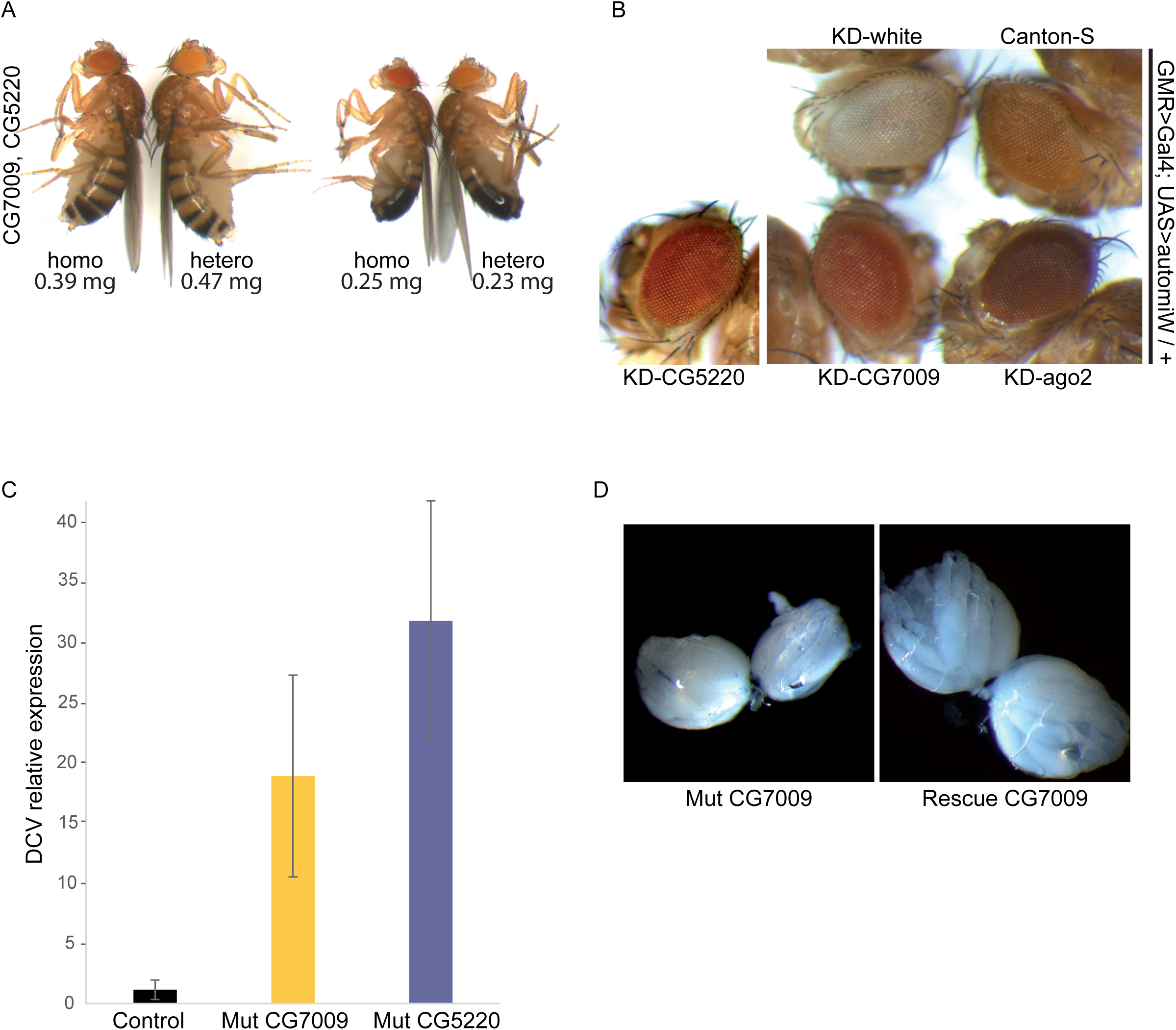
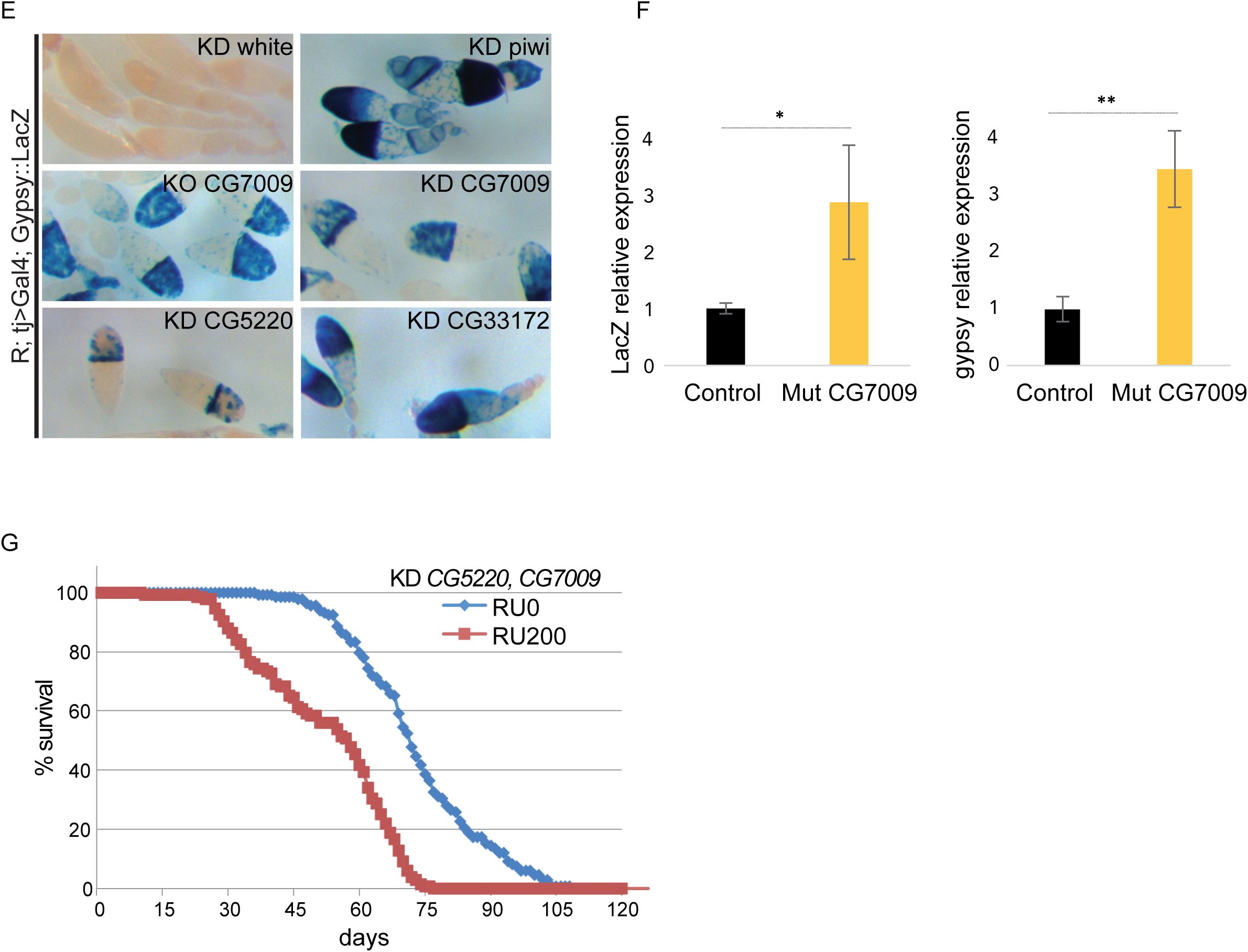
CG7009 and CG5220 affect small RNA silencing pathways and life span. **A**, Homozygous *CG7009, CG5220* double mutant flies display reduced adult weight and size. Images of adult females and males *CG7009, CG5220* homozygous double mutants (homo) compared to heterozygous double mutants (hetero). Below the images is indicated the average weight for flies in milligrams (mg) calculated for 3-day-old flies measured on precision balance. *p*-value<0,001 in a Student’s T-test. **B**, CG7009 and CG5220 modulate the Ago2-dependent gene silencing in somatic tissues. The UAS>*automiW* construct is a sensor derived from *automiG* where two miRNAs were reprogrammed to target the *white* gene (47). KD indicates eye-specific GMR-Gal4/UAS-RNAi-mediated inactivation of the respective genes (*white*, *CG7009*, *CG5220* or *Ago2*). Canton-S *was* used as control for eye color. Darker eye coloration than Canton-S (top right) indicates that the sensor is failing to inactivate the *white* gene through Ago2-dependent silencing. Flies images were taken at the same age (5 days old). **C**, The siRNA-dependent viral defence is compromised in *CG7009* and *CG5220* mutants. RT-qPCR using *Drosophila* C Virus (DCV) specific primers three days after injection with DCV solution or solution free of DCV as control (not shown) in heterozygous *CG7009^e02001^* mutants (control) or homozygous *CG7009^e02001^* (CG7009) and *CG5220^K>A^* (CG5220) mutants. Represented is the relative DCV expression to Rp49. Error bars represent the mean ± s.d (standard deviation) between biological replicates (n=2 replicates where n is a mix of 2 to 3 flies). **D**, *CG7009* mutation is associated with ovarian size reduction. The images show representative examples of ovaries from a 4 days old fertilized female raised on fresh yeast of trans-heterozygous *CG7009^e02001^/ Def9487* mutant (Mut CG7009) and a genetic rescue ovaries of *CG7009 (BAC)/ +; CG7009^e02001^ / Def9487* mutant (Rescue CG7009); n>15 for each genotype. WT ovaries (not shown) were similar to rescued ovaries of *CG7009* mutant; Mut: mutant. **E**, CG7009 and CG5220 are involved in *gypsy* TE-repression in *Drosophila* ovaries. *Gypsy::lacZ* sensor is silenced in follicle cells using tj>Gal4-mediated expression of an UAS-RNAi line (KD) against the *white* gene (KD control, R; tj>Gal4/ +; Gypsy::lacZ/UAS-*white*-RNAi), no blue coloration = no β-Gal staining). *Gypsy* silencing is disrupted using *piwi* KD (positive control: blue coloration = positive β-Gal staining) and after KD against the indicated genes. The *Gypsy::lacZ* sensor is also de-repressed in null *CG7009* homozygous mutant (KO). KD: knock down, KO: knock out. **F**, RT-qPCR using *gyspy* or *lacZ* specific primers on ovaries from flies KD for *white* (control) or *CG7009* expressing the *Gypsy::lacZ* sensor (tj>Gal4/ +; Gypsy::lacZ/UAS-RNAi) as described in Figure 2E). Error bars represent the mean ± s.d (standard deviation) between three independent biological replicates. **G**, *CG7009, CG5220* double KD flies display reduced life span. Survival curves of males double KD for *CG5220*, *CG7009* with (RU200) or without (RU0) RU486-mediated RNAi transgene induction. Constitutive expression (RU200) of *CG5220*, *CG7009* KD transgenes was induced by RU486 exposure (20 mg/ml during adulthood). The curves represent the average values of at least three biological replicates of 10 flies per experiment.

### CG7009 and CG5220 contribute to efficient miRNA Ago2-mediated RNA silencing *in vivo*

To address whether CG7009 affected small RNA silencing pathways *in vivo*, we expressed the *automiW* sensor, which is based on the knockdown of the *white* gene by means of *white*-targeting miRNAs in the developing eye (47). Combining this sensor construct with RNAi-mediated knockdown of CG7009 or CG5220, we observed increased eye coloration when compared to controls (Figure 2B). This observation indicated that Ago2-dependent silencing or miRNAs biogenesis of this reporter was affected non-redundantly when *CG7009* or *CG5220* were post-transcriptionally down-regulated. These results implicated both genes in proper functioning of either miRNA biogenesis and/or Ago2-dependent and small interfering RNA-mediated silencing *in vivo*.

### siRNA-mediated RNA silencing is impaired in *CG7009* and *CG5220* mutant flies

As small interfering RNA-mediated silencing is required for viral defence in *Drosophila (48)*, we tested whether viral defence was impaired in *CG7009* or *CG5220^K>A^* mutant flies. To this end, purified *Drosophila* C virus (DCV) was injected into the thorax of adult flies and the viral load was monitored by qRT-PCR 4 days after infection. The results of these experiments showed that flies lacking CG7009 or CG5220 function were significantly more sensitive to DCV infection when compared to control flies (Figure 2C). The results indicated that *CG7009* and *CG5220^K>A^* mutants failed to initiate or maintain a proper response to viral infection, which, together with the results of our genetic screen using *automiG* for miRNA biogenesis and Ago2-dependent silencing of gene expression (Figure 1B and 2B) and the siRNA activity reporter assays (Supplementary Figure S1B), supported the notion that both gene products were required for efficient Ago2-dependent and small interfering RNA-mediated silencing activities in *Drosophila*.

### piRNA-mediated RNA silencing is affected in *CG7009* and *CG5220* mutant flies

During the characterization of the *CG7009* mutants, we noticed that females ovaries were reduced in size when compared to BAC-rescued control flies (Figure 2D). This ovarian phenotype was mild but similar phenotypes were previously described in several mutants for genes required in the piRNA pathway (49). Although the genetic screen was specifically designed to identify genes involved in miRNA biogenesis or Ago2-mediated silencing pathways, we tested whether CG7009 and CG5220 function also affected transposable element (TE) silencing, especially through the somatic piRNA pathway. To this end, the activity of a somatic piRNA-mediated silencing reporter (50) was monitored in adult ovaries. This reporter faithfully recapitulates the expression of the retro-transposon *gypsy* in ovarian follicle cells, in which abundant somatic piRNAs are produced in defence against mobile elements (50). Remarkably, piRNA-mediated silencing of this reporter was de-repressed upon somatic follicle cell-specific knockdown of *CG5220* and *CG7009* expression (Figure 2 E-F). Furthermore, expression of endogenous *gypsy* mRNA was elevated in *CG7009* mutants (Figure 2F). Taken together, these results suggested that both genes are involved in TE silencing mediated by the somatic piRNA pathway in *Drosophila*.

### Mutation in CG7009 and CG5220 affects life span

Although a size reduction of *CG7009*, *CG5220* double mutant adult flies could be observed (Figure 2A), no other obvious mutant phenotypes affecting flies morphology could be observed. Importantly, *CG7009^e02001^*,*CG5220^K>A^* double mutant flies displayed reduced lifespan when compared to controls (Figure 2G). Mutant flies lived, on average, about 25 days shorter than controls. Furthermore, homozygous *CG7009^e02001^* mutant flies as well as *CG7009^e02001^*,*CG5220^K>A^* double mutants appeared sluggish and less active displaying general locomotion defects (not shown).

### CG7009 and CG5220 are Nm-MTases acting on tRNAs

To test whether CG7009 is an Nm-MTase, recombinant proteins were expressed and purified from *E. coli*. *In vitro* methylation assays using *in vitro*-synthesized *Drosophila* tRNA^Phe^ did not reveal activity of recombinant CG7009 protein. In order to ascertain the predicted catalytic activities of CG7009 and CG5220, we analyzed the Nm methylation status of *Drosophila* tRNA^Phe^, which is a substrate of TRM7 in yeast, using control, *CG7009^e02001^* and *CG5220^K>A^* mutants flies. We performed sequence-specific purification of tRNA^Phe^ using biotinylated DNA oligonucleotides coupled to streptavidin matrices followed by RNase digestion and MALDI-TOF mass spectrometry. RNase A has a preference for hydrolysis at pyrimidine residues, while RNase T1 is strictly guanosine-specific. Because Nm at a given nucleotide position (*n*) protects the adjacent 3’-phosphodiester bond to the neighbouring nucleotide (position *n+1*) against nuclease attacks, various specific digestion products of *Drosophila* tRNA^Phe^ can be expected as a result of RNase A or RNase T1 activities. In addition, according to the reported modification profile of *Drosophila* tRNA^Phe (51, 52)^, which includes Nm at C_32_ and G_34_, very specific RNA fragments were thus predicted (Figure 3A and Supplementary Figure S3).

**Figure 3.**
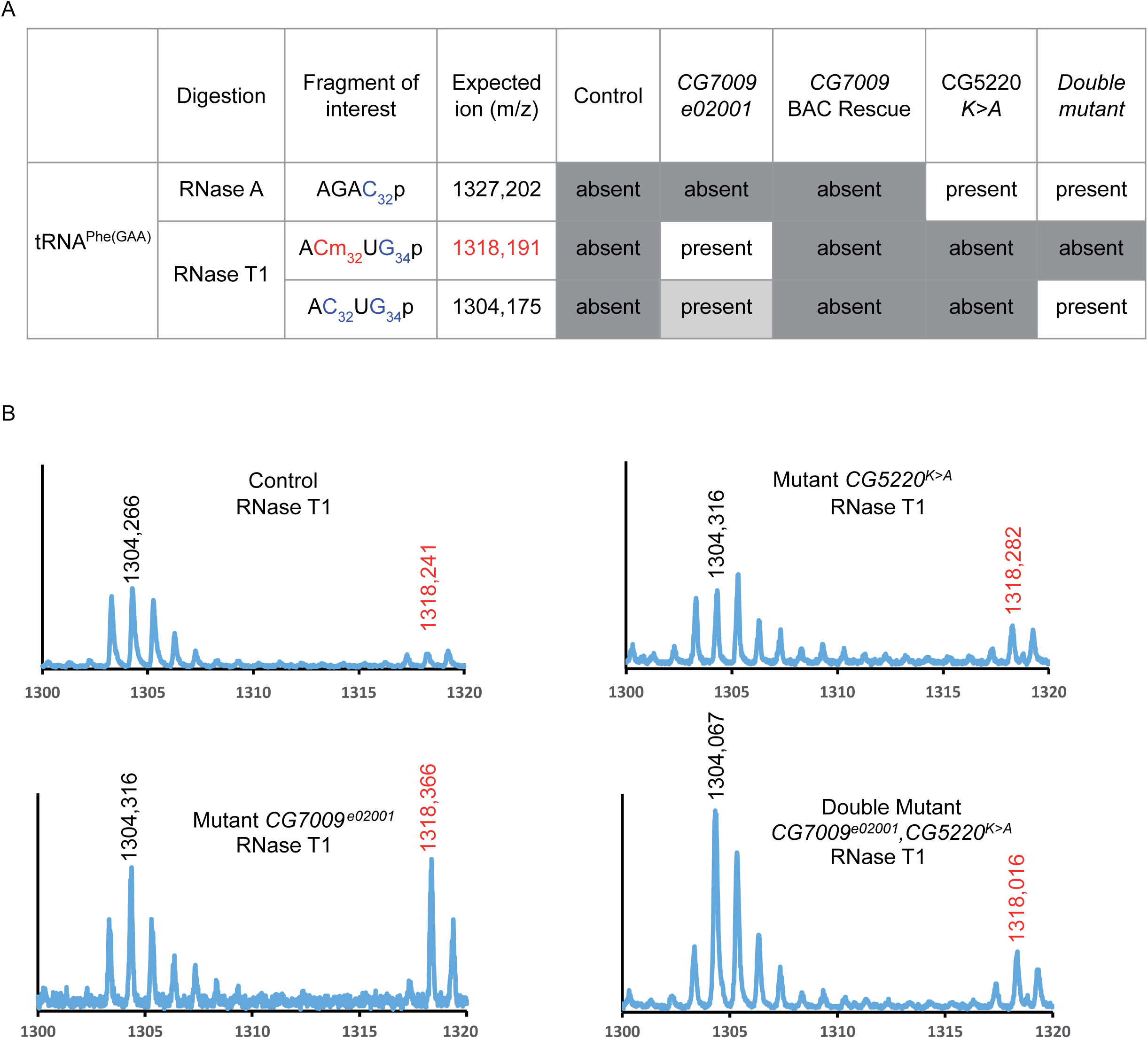
CG7009 & CG5220 are TRM7-like tRNA Nm MTases in *Drosophila*. **A**, Presence (present) or decrease (absent) of Nm-containing RNA fragments and their sizes (m/z in Daltons) upon RNase A or T1 digestion of tRNA^Phe(GAA)^ extracted from *Drosophila* adult heterozygous *CG7009^e02001^* mutants (Control), homozygous *CG7009^e02001^* mutants (CG7009*^e02001^*), homozygous *CG7009* mutants rescue BAC (CG7009 BAC Rescue), homozygous *CG5220^K>A^* mutants (CG5220 *K>A*), and double homozygous mutants *CG7009*^e02001^,*CG5220^K>A^* (Double mutant). **B**, MALDI TOF-MS spectrum of fragments resulting from RNase T1 digestion of tRNA^Phe-(GAA)^ originating from heterozygous adult *CG7009^e02001^* mutants (Control), homozygous *CG7009^e02001^* mutants (Mutant *CG7009^e02001^*), homozygous *CG5220^K>A^* mutants (Mutant CG5220*^K>A^*) and double homozygous mutants *CG7009^e02001^*,*CG5220^K>A^* (Double Mutant *CG7009^e02001^*,*CG5220^K>A^*) as indicated. Relevant peaks are identified by their m/z values (X abscise). **C**, RiboMethSeq analysis of tRNA^Phe(GAA)^ Nm modifications at positions C_32_ and G_34_. Alkaline fragmentation-based RiboMethSeq protocol was performed on total RNAs extracted from whole heterozygous flies (Control) or homozygous mutants for *CG7009^e02001^* (CG7009*^e02001^*) and Rescue *CG7009^e02001^* (BAC) as indicated. Normalized cleavage efficiency, calculated from cumulated 5’-end and 3’-end coverage, is shown for the +/− 5 neighboring nucleotides. The positions of interest (C_32_ and G_34_) in tRNA^Phe(GAA)^ are pointed with red arrows. Protection against cleavage is indicated (+) protected, (−) not protected. Protection at Cm_32_ in control flies is only moderate, indicating incomplete tRNA methylation (+low). Higher methylation levels at this position are observed in *CG7009* homozygous mutants. **D**, Methylation scores (MethScore) for 5 tRNA 2’-O-methylated positions showing altered methylation in *CG5220* and/or *CG7009* indicated mutants. MethScore (Score C), representing the level of ribose methylation was calculated from protection profiles. Data are shown for positions 32 and 34 in different *D. melanogaster* tRNAs in control flies heterozygous for *CG7009^e02001^*, homozygous mutant *CG5220^K>A^*, two independent genetic background mutants for *CG7009* (homozygous *CG7009^e02001^* or mutant trans-heterozygous *CG7009^e02001^*/*Def3340*), double homozygous mutant *CG7009^e02001^*,*CG5220^K>A^* and rescue BAC *CG7009^e02001^*/*Def3340* flies.

First, we determined fragments that were obtained after RNase A hydrolysis of tRNA^Phe^, which should provide information on the Nm-modification status at C_32_. MALDI-TOF analysis revealed almost no fragment of 1327.2 Da (AGAC_32_p fragment) in control flies indicating that C_32_ was modified with Nm thereby blocking RNase A activity at this position in tRNA^Phe^ from wild type and rescue flies (Figure 3A and Supplementary Figure S4A, B). This fragment increased significantly in *CG5220^K>A^* mutants suggesting loss of protection from RNase A activity in animals without CG5220. Interestingly, the increase in RNase A-mediated tRNA^Phe^ fragmentation observed in *CG5220^K>A^* mutants could only be moderately observed when using tRNA^Phe^ from *CG7009^e02001^,CG5220^K>A^* double mutant flies (Figure 3A and Supplementary Figure S4A) indicating that C_32_ protection from RNase A is largely independent of CG7009. In support of this notion, the *CG7009^e02001^* mutation alone did not affect the RNAse A digestion profiles when compared to control (Figure 3A and Supplementary Figure S4A) or BAC rescue *CG7009^e02001^* flies (Supplementary Figure S4B). These results indicated that CG5220, but not CG7009, harbors an activity that protects tRNA^Phe^ at C_32_ against RNase A digest, therefore making CG5220 the main candidate for an Nm-MTase at this position in *Drosophila*.

Next, we obtained RNase T1 digestion profiles to deduce the G_34_ modification status of tRNA^Phe^ in both control and mutant flies. MALDI-TOF analysis showed a ACm_32_UG_34_p fragment (1318,1 Da) that could not be detected in wild type flies indicating that G_34_ was modified with Nm thereby blocking RNase T1 activity at this position in wild type tRNA^Phe^ (Figure 3A, B). This fragment increased significantly in *CG7009^e02001^* mutants suggesting loss of protection from RNase T1 activity in animals without CG7009. The RNase T1 digestion profiles from controls and *CG5220^K>A^* mutant flies were comparable (Figure 3A, B and Supplementary Figure S4B) indicating that CG7009 but not CG5220 is implicated in protecting G_34_ from RNase T1 digestion in tRNA^Phe^. Finally, digest of tRNA^Phe^ from *CG7009^e02001^, CG5220^K>A^* double mutant flies with RNase T1 produced a fragment (AC_32_UG_34_p) that was completely unmodified (1304 Da) suggesting that CG5220 and CG7009 are the responsible Nm-MTase activities that modify C_32_ and G_34_ in tRNA^Phe^, respectively (Figure 3A, B). Notably, Nm deposition at G_34_ was not affected in *CG5220^K>A^* mutants (Figure 3A, B) indicating that CG5220-mediated modification at C_32_ does not influence the deposition at neighboring G_34_ . In contrast, loss of Nm at G_34_ in *CG7009^e02001^* mutants moderately affected Nm C_32_ deposition (Figure 3A, B, 1304 Da). These results indicated that while Nm at C_32_ was not a prerequisite for Nm at G_34_, Nm at G_34_ was probably required for the correct formation of methylation at C_32_ or for the stability of the methylated tRNA.

Collectively, these data demonstrated that genetic mutation of two candidate Nm-MTases in *Drosophila* resulted in the reciprocal loss of two conserved ACL modifications in tRNA^Phe^ strongly suggesting that CG5220 and CG7009 are indeed functional methyltransferases responsible for the deposition of Nm at C and G of tRNA^Phe^, respectively. Interestingly, our results also suggest that *Drosophila melanogaster*, and most likely all other *Drosophila* species, evolved two distinct TRM7 family members to methylate the ACL on substrate tRNAs (Figure 1D).

### Methylation specificity of both MTases depends on nucleotide position

To obtain a comprehensive picture of the Nm specificity for CG7009 and CG5220 *in vivo*, we performed RiboMethSeq analysis on *Drosophila* tRNAs. RiboMethSeq allows RNA-wide Nm detection based on random RNA fragmentation by alkaline hydrolysis followed by library preparation and sequencing (53, 54). The presence or absence of Nm can be appreciated from characteristic coverage profiles of the 5′-/ 3′-ends of cDNAs. Since Nm residues protect the adjacent 3’-phosphodiester bond to the neighbouring nucleotide from hydrolysis, a gap in the coverage at the *n+1* position indicates the presence of a 2’-O-methylated nucleotide at position *n*.

When analyzing the 2’-O-methylation status at position 34 for tRNA^Phe^ in control individuals, reads at position 35 (equals *n+1)* were under-represented in regard to their direct nucleotide neighbors (Figure 3C). This demonstrated that, Nm was present at G_34_ in *Drosophila* tRNA^Phe^ as previously reported (51) and as shown by MALDI-TOF MS analysis (Figure 3A). Similarly, RiboMethSeq profile analysis of *CG5220^K>A^* mutants indicated G_34_ to be methylated (Supplementary Figure S5A). The presence of Nm at G_34_ in *CG5220^K>A^* mutant confirmed that CG5220 was not involved in the formation of ribose methylation at this position. On the contrary, in two different *CG7009* mutants, as well as in *CG7009^e02001^*, *CG5220^K>A^* double mutants, protection against hydrolysis at position 35 was totally abolished when compared to the control heterozygote profile (Figure 3C and Supplementary Figure S5A) confirming that CG7009 is the Nm-MTase for G_34_ of tRNA^Phe^ in *Drosophila* and that CG5220 alone is not able to methylate this position. Importantly, the expression of an additional gene copy of CG7009 in the *CG7009^e02001^* mutant background (Rescue CG7009 BAC) rescued the loss protection against hydrolysis at G_34_ of tRNA^Phe^ (Figure 3C and Supplementary Figure S5C). In addition, RiboMethSeq analysis was performed for position 33 of tRNA^Phe^ (*n+1* to the expected Nm at C_32_ ((51) and Figure 3A), which confirmed that CG5520, but not CG7009, was responsible for ribose methylation at position 32 on tRNA^Phe^ (Figure 3C, D and Supplementary Figure S5A, C).

Furthermore, RiboMethSeq analysis also identified other tRNAs potentially methylated by CG7009 and CG5220, some of which were already known as substrates of TRM7 orthologs in other species. For instance, alike tRNA^Phe^, we found CG7009-dependent methylation in tRNA^Trp^ at position C_34_ as well as CG5520-dependent Nm at position C_32_ (Figure 3D). Strikingly, the methylated nucleotide at position 34 in tRNA^Trp^ of *Drosophila* is a cytosine, like in humans and in yeast (1, 35, 38, 39). Importantly the RiboMethSeq profiles clearly showed that CG7009 (and not CG5220) methylated this position (Figure 3D) indicating that CG7009 can deposit Nm on G and C nucleotides. The same observation was made for CG7009-mediated methylation of C_34_ in tRNA^Leu(CAA)^, in agreement with previous data showing that FTSJ1 was responsible for depositing Nm at f5C_34_/hm5C_34_ in human tRNA^Leu(CAA)^ (34) and Supplementary Figure S5B, C). In addition, we identified previously unknown Nm-MTase substrate tRNAs. We found CG5220-dependent methylation of tRNA^Gln^ and tRNA^Glu^ at position C_32_ (Figure 3D and Supplementary Figure S5C). 2’-O-methylated C_32_ in tRNA^Glu(UUC)^ had been previously reported in *Drosophila* (1, 55). Interestingly, cytosine 32 was also reported to be 2’-O-methylated in human tRNA^Gln^ by a yet unidentified enzyme (1). Our data thus suggest that the human ortholog of CG5220, FTSJ1, may be the Nm-MTase responsible for the modification at this position.

Altogether, detailed RiboMethSeq analysis confirmed the MALDI-TOF MS results (Figure 3A, B) demonstrating that CG5220 is specialized for depositing Nm at C_32_ nucleotides while CG7009 is responsible for modifying the wobble position. Furthermore, the discovery of additional tRNA substrates (Supplementary Figure S5C) for both Nm-MTases suggested that their respective specificity is dependent on the position rather than on the nature of nucleotide (C, U or G).

### CG33172 is part of the Nm-MTase complex

The yeast TRM7 associates with two distinct proteins that are required for its catalytic activity (35, 36). Deposition of Nm at C_32_ by TRM7 is supported by binding to TRM732 while the interaction with TRM734 is necessary for addition of Nm at position 34. THADA and WDR6 are the orthologs of TRM732 and TRM734 in humans, respectively, and the interactions with FTSJ1 are conserved (39). In *Drosophila*, CG15618, also known as DmTHADA (56), is the potential ortholog of TRM732 and THADA, while CG33172 is the putative orthologue of TRM734 and WDR6 (Supplementary Figure S6A). Importantly, CG33172, TRM734 and WDR6 are members of the WD40-repeat-containing domain superfamily that contains also the human protein WDR4, another tRNA-MTase cofactor involved, as FTSJ1, in neurodevelopmental disorders (57, 58).

The use of the *automiW* sensor combined with dsRNA-mediated knockdown of *CG15618* and *CG33172* in the *Drosophila* eye recapitulated the Ago2-mediated small RNA silencing failure observed in *CG7009* and *CG5220* mutants (Figure 4A). Interestingly, dsRNA-mediated knockdown of *CG33172* using the *gypsy-lacZ* sensor also recapitulated the somatic piRNA silencing failure observed in both *CG7009* and *CG5220* mutants (Figure 2E), indicating genetic interactions between CG7009/ CG5220-mediated functions and these gene products.

**Figure 4.**
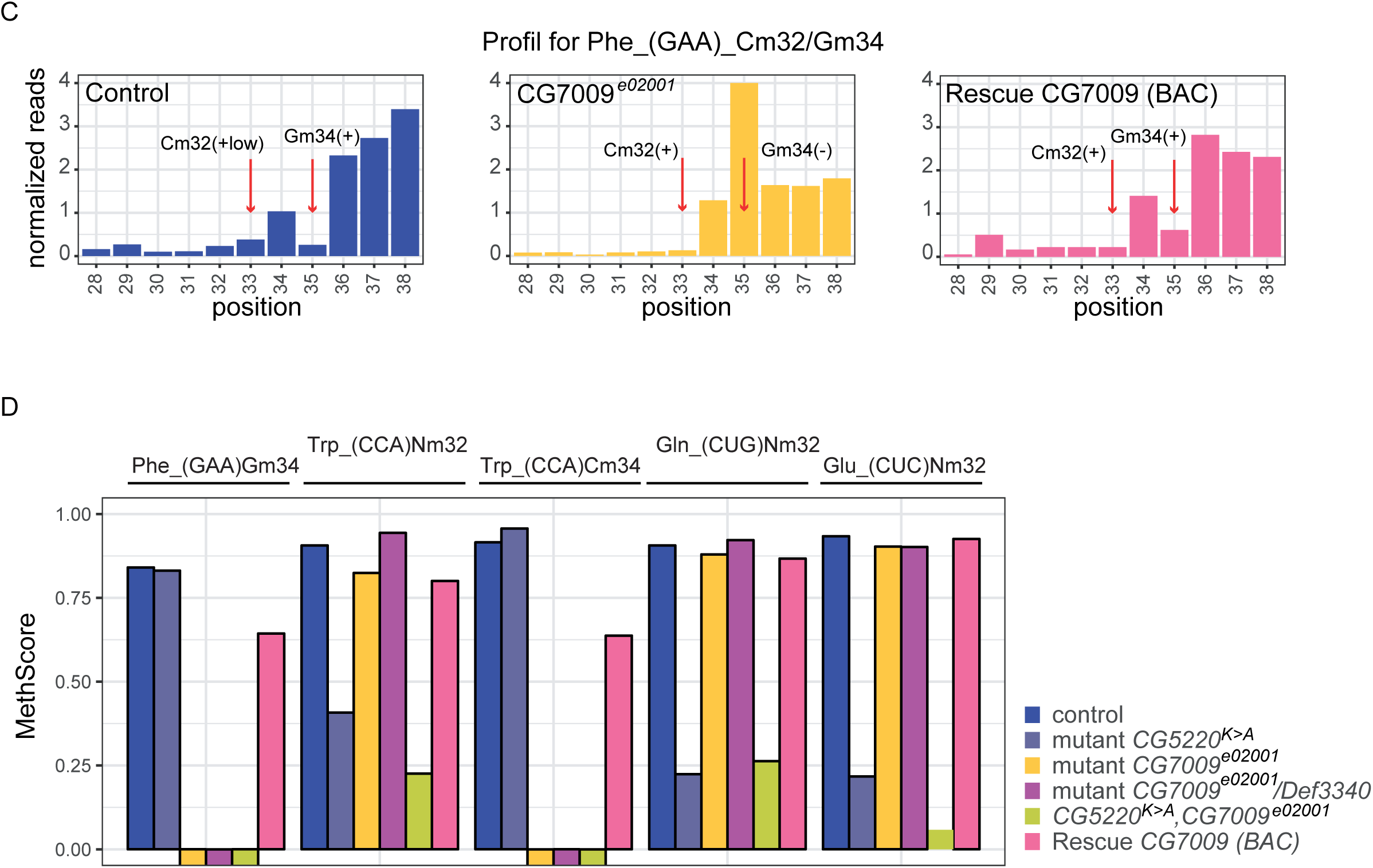
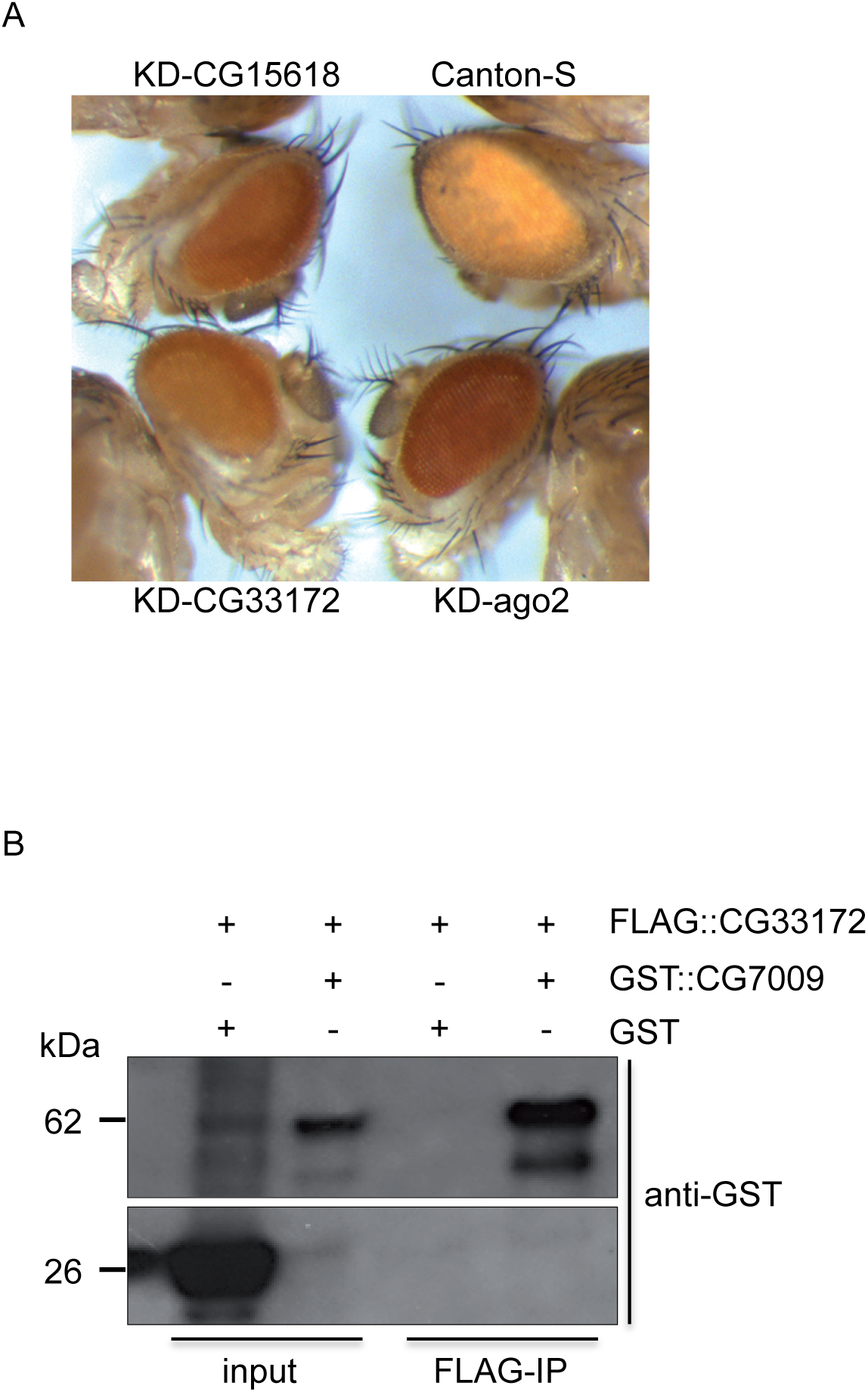
CG33172 is a partner of the Nm-MTase complex in *Drosophila*. **A**, CG33172 and CG15618 modulate Ago2-dependent silencing in adults flies. *CG33172* and *CG15618* were knocked down by using UAS-RNAi (KD) lines and eye specific GMR-Gal4, w-driver (indicated as KD), as in Figure 2B. Canton-S (wild type w+) and *Ago2* KD were used as controls. A darker eye coloration than Canton-expressing *automiG* lines (top right) indicates that the miRNAs of the sensor are failing to inactivate the *white* gene through Ago2-dependant silencing. **B**, CG33172 interacts *in vitro* with CG7009. Co-immunoprecipitation of tagged CG7009 and CG33172 recombinant proteins co-expressed in bacteria. Anti-GST WB on co-expressed GST::CG7009 and FLAG::CG33172 inputs and after FLAG-IP; Lower panel, Anti-GST WB reveals a GST “alone” signal in the co-expressed GST and FLAG::CG33172. Inputs correspond to 10% of 10 µg of protein eluates. The expected proteins sizes are GST (26 kDa) and GST::CG7009 (62 kDa). WB - Western blot; kDa - kilodaltons.

In order to test for physical interactions between CG7009, CG15618 and CG33172, we cloned FLAG-tagged CG15618 and CG33172 with the aim of co-expressing these proteins along with GST::CG7009 in bacteria. While co-expression of FLAG::CG15618 was technically challenging due to the size of this protein (197 kDa), FLAG::CG33172 could be expressed and immunoprecipitated using anti-FLAG antibodies. The precipitate was tested for the presence of GST::CG7009 by using western blotting and anti-GST antibodies. The results showed that FLAG::CG33172 co-precipitated with GST-CG7009 but not GST alone suggesting a direct interaction between these two proteins (Figure 4B). Collectively, these observations suggested the existence of an Nm-MTase complex containing CG7009 and at least one accessory protein, CG33172, which might be required for depositing Nm at position 34 on selected tRNAs.

### Nm limits endonucleolytic cleavage of tRNA^Phe^

We next addressed the mechanisms underlying the defects in the Ago2-mediated RNA silencing activity and life span observed in *CG7009* mutant flies. It has been reported that loss of m^5^C from specific tRNAs resulted in increased tRNA fragmentation in *Drosophila*. Interestingly, it was proposed that tRNA fragments could affect small RNA silencing pathways through binding to Dicer and Argonaute proteins thereby reducing their activity (59). In addition, during the preparation of this manuscript, a study showed that Nm protected tRNA^Met(CAT)^ from endonucleolytic cleavage by stress-induced angiogenin in human cell (9).

We therefore tested if lack of Nm at positions 32 and 34 of *Drosophila* tRNAs affected their endonucleolytic cleavage during stress conditions. A heterozygous *CG7009^e02001^* mutant (control), a trans-heterozygous mutant for CG7009 (*CG7009^e02001^*/*Def*) and the rescue line for CG7009 (BAC) were analyzed by northern blotting with a specific probe complementary to the 5’-end of tRNA^Phe^ before and after heat shock. Two clear hybridization signals were observed, corresponding to mature tRNAs (70-75 nt) and tRNA fragments (30-35 nt, Figure 5A). tRNA fragmentation increased significantly in *CG7009^e02001^*/*Def* mutants. Importantly, increased tRNA fragmentation was rescued in BAC flies (Figure 5A), demonstrating that CG7009 function affected tRNA fragmentation of tRNA^Phe^. Of note, global steady state levels of mature tRNAs were not affected in *CG7009^e02001^*/*Def* mutants (Figure 5A), suggesting limited translational defects in flies without functional CG7009. Interestingly, we observed no stress-dependent effects on tRNA fragmentation in *CG7009^e02001^*/*Def* or other *CG7009* mutant combinations (Figure 5A, B). Furthermore, we did not observe tRNA^Phe^ fragmentation changes in *CG5220^K>A^* mutants (Figure 5B). Thus, these results indicated that Nm at position G_34_ limits fragmentation of tRNA^Phe^ suggesting that 3’ terminal Cm stabilizes the observed tRNA fragments in *CG7009* mutants.

**Figure 5.**
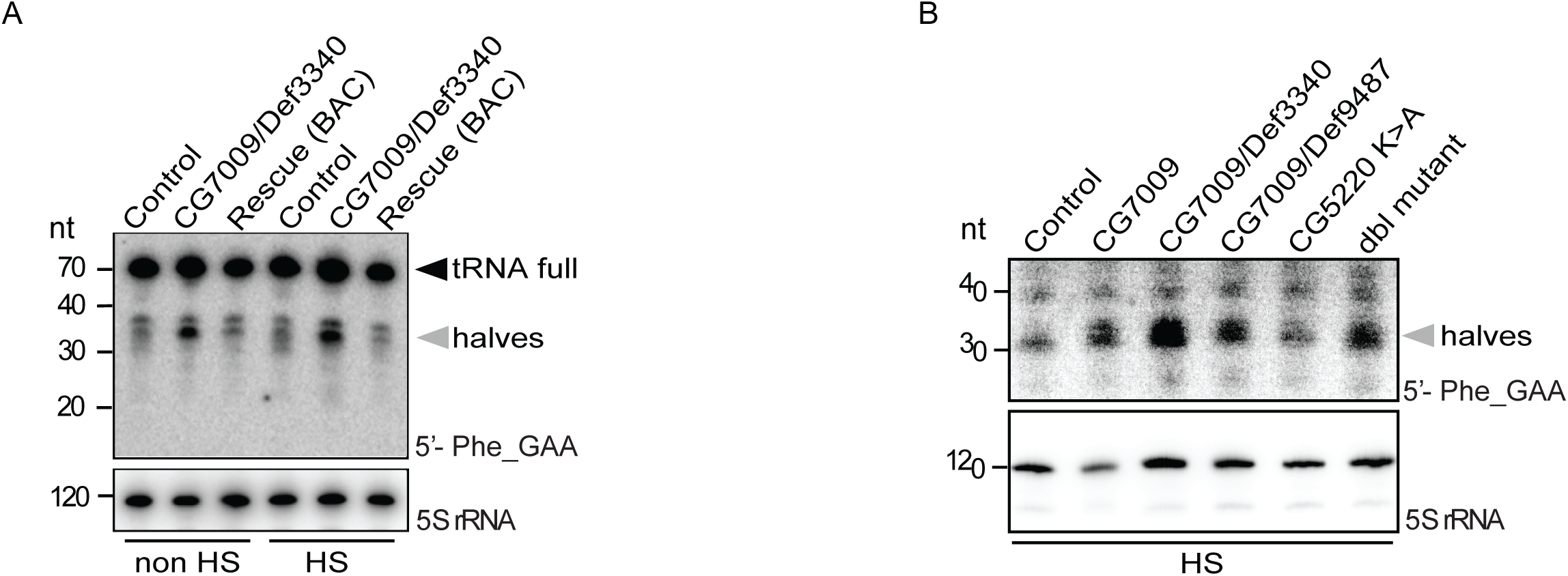
Nm34 limits tRNA endonucleolytic cleavage in adult flies. **A**, Northern blot characterization of 5’-tRNA^Phe(GAA)^-derived tRFs. Northern blot on total RNAs extracted from whole heterozygous *CG7009^e02001^* flies (control), transheterozygous for *CG7009^e02001^* / *Def3340* or rescued mutants for *CG7009* (rescue BAC) as indicated using a 5’-tRNA^Phe(GAA)^ specific probe and a 5S rRNA probe as loading and transfer control. Mature tRNA^Phe^ size is 73 nt (full length). 5’-tRNA^Phe^-derived tRNA fragments (5’-tRF^Phe^) were detected at ∼35 nt (halves). The same experiment was performed on heat-shocked flies (1 hours at 37°C in incubator), RNAs were extracted after 5 hours at 25°C (indicated as HS, Heat Shock). nt: nucleotide. **B**, The same experiment as in Figure 5A above was performed on heat-shocked flies (1 hours at 37°C in incubator), RNAs were extracted after 5 hours of flies recovery at 25°C (indicated as HS, heat shock) with additional indicated genotypes. nt: nucleotide; dbl mutant: Double Mutant *CG7009^e02001^*,*CG5220^K>A^*.

## Discussion

Surprisingly, while performing an RNAi genome-wide screen for modulators of the Ago2-dependent small RNA silencing in *Drosophila*, we identified a previously uncharacterized gene (*CG7009*) with sequence homology to Nm-MTases of the TRM7/FTSJ1 subfamily of MTase proteins. Through sequence analysis, we identified an additional gene (*CG5220*) in *Drosophila*, which shares considerable sequence homology with CG7009. By characterising the molecular function of these predicted Nm-MTases, we demonstrate that both genes encode for RNA methyltransferases depositing Nm on tRNAs and displaying specialized activity at two distinct positions in the ACL. These findings reveal that, in contrast to both yeast and humans, which encode only one Nm-MTase gene capable of methylating the ACL, *Drosophila melanogaster* has evolved two enzymes that methylate two nucleotides in the ACL of conserved tRNA targets. Interestingly, it appears that all *Drosophila* species evolved and maintained two closely related TRM7/FTSJ1 paralogs. This peculiarity of *Drosophila* species makes us proposing that Nm in the ACL is deposited in sequential order either by one enzyme (TRM7 or FTSJ1 in yeast or human, respectively) or by two enzymes in *Drosophila* species. We suggest that Nm at the wobble position is a prerequisite for proper deposition or maintenance of Cm at position 32. Importantly, our analysis also reports novel substrates of the TRM7 subfamily of Nm MTases.

However, and in agreement with studies on various RNA modification enzymes in other organisms, Nm-MTases in *Drosophila* are not required for viability nor fertility. Nevertheless, Nm-MTases mutants displayed reduced life span and behavioural phenotypes manifested as general mobility defects (not shown). Abolishing the catalytic function of both genes in the same animal did not reveal other obvious mutant phenotypes with the notable exception of a reduction in size and weight, highlighting a potential role of these Nm-MTases in specific, but not general, translational control as previously reported for *trm7* mutant yeast (38, 60).

Nm modifications in the ACL of specific tRNAs can affect translational efficiency and fidelity (61). Consistently, Nm deposition in mRNA also affected translation through interference with tRNA decoding efficiency and thus could potentially rewire the genetic code (62, 63). Interestingly, it was recently proposed that TRM7 methylates substrates different than tRNAs including mRNAs in yeast (16, 17) suggesting that TRM7 family members can act as multi-substrate Nm-MTases, which modulate translation by methylating mRNA codons and tRNA anticodons. Importantly, loss of Nm at tRNA positions 32 and 34 in *trm7* mutant yeast affected translation rates and, consecutively, cell growth (36, 38). Thus, the observed reduction in size and mass (Figure 2A) in flies without TRM7 family members supports the hypothesis that *CG5220* and *CG7009* mutations affect translational efficiency in *Drosophila*.

Importantly, the lack of Nm at the wobble position in *CG7009* mutants facilitated tRNA fragmentation. Transfer RNA fragmentation is a conserved response to various stress conditions with functions ranging from protein synthesis, to apoptosis and the modulation of small non-coding RNA pathways (64–67). The influence of internal Nm modifications on tRNA stability has only been described very recently (9). In support of the notion that Nm in tRNAs modulates their stability, we found that Nm_34_ is protective against tRNA fragmentation in the ACL. However, in contrast to the Nm-mediated effect on tRNA cleavage in *CG7009* mutants, the absence of Nm_32_ in *CG5220* mutants did not change tRNA fragmentation patterns. Since Hen1-mediated deposition of Nm at RNA 3’-termini (13, 68) stabilizes small RNAs in various organisms, we propose that tRFs produced by ACL cleavage in *CG7009* mutants may be stabilized through the existence of 3’ terminal Nm_32_ deposited by CG5220, explaining the abundance and apparent stability of tRFs in *CG7009* single mutant in contrast to *CG5220* or *CG7009*, *CG5220* double mutants. To the best of our knowledge this is the first report that involves a TRM7 family protein in tRNA fragmentation.

Importantly, loss of function of either CG7009 or both Nm-MTases impaired Ago2- as well as Piwi-dependent small RNA silencing pathways *in vivo*. Furthermore, DCV infection assays in adult flies confirmed a function for CG7009 and CG5220 as regulators of siRNA pathway-mediated virus control. These results indicated that Nm-MTases function affects small RNA silencing pathways in a pleiotropic fashion and placed these genes upstream of small RNA biogenesis and function. Indeed, RNA sequencing analysis suggested that the manifestation of these phenotypes could partially be due to the transcriptional downregulation of Ago2 mRNA in *CG7009* mutant flies or after knockdown of CG7009 in S2 cells.

On the other hand, tRNA fragments can associate with Dicer, Argonaute and Piwi proteins (59, 69–71). One potential consequence of such interactions could be a reduction in the capacity of small RNA pathway components to process or bind to canonical RNA substrates. Indeed, tRF-mediated titration of proteins has been reported (72–74). It remains to be tested whether also the newly identified Nm-MTase substrates (*i.e.*, tRNA-Trp, Leu, Gln, Glu) accumulate tRFs in *CG7009* and/or *CG5220* mutant tissues, which would support the notion that loss of Nm in tRNAs is causative for decreased tRNA stability and thereby could also contribute to the failure of Ago2/Dcr2-dependent small RNA pathways.

Finally, our study identifies CG33172 as a binding partner of CG7009. Interestingly, the yeast ortholog of CG33172, TRM734, was reported to control the steady state levels of TE as does TRM7 (75). Here, we have shown that the *Drosophila* orthologs of TRM734 and TRM7 (CG331772 and CG7009, CG5220, respectively) also affect TE silencing pathways. Importantly, mutations in human WDR4 impaired tRNA m^7^G_46_ methylation and caused microcephalic primordial dwarfism (58). CG33172 belongs to the WD-Repeat (WDR) family of proteins and is the ortholog of human WDR6. Interestingly, both WDR6 and FTSJ1 were identified as principal human host restriction factors against *vaccinia* virus indicating that this Nm-MTase complex functions at the interface of host-virus interactions (76). In support of the notion that Nm modifications modulate mobile element control, human Nm-MTase FTSJ3 can be hijacked by HIV-1 to methylate viral mRNAs resulting in avoidance of being sensed by the host innate immune system (17).

Our study thus strongly supports the emerging notion that an important biological impact of Nm-MTase activity is mobile element control affecting TEs and viruses. Importantly, our results in *Drosophila* also indicate that the molecular machinery necessary to deposit Nm in tRNAs and the associated physiological importance are conserved throughout evolution.

In summary, this study provides a comprehensive *in vivo* characterization of two Nm MTases enzymes and associated functions in *Drosophila melanogaster*, demonstrating the crucial importance of the TRM7 family Nm MTase enzymes in controlling life span and small non-coding RNA silencing functions. The peculiarity of *Drosophila* species that evolved two TRM7 genes products with specialized activity in specific tRNAs showed that Nm in the ACL is deposited sequentially. The Nm at wobble position of tRNA^Phe^ is a prerequisite for proper Nm modification at position 32.

## Methods

### Automig DRSC 2.0 genome-wide screening library

We screened 22.490 dsRNA of 400 bp length on average allowing the inactivation of 94,3 % of the annotated *Drosophila* genes, including predicted genes (library version DRSC 2.0). About 13.900 genes are represented in the collection (∼66 assay plates), on average of one to two dsRNAs per gene. More information about the DRSC Genome-wide RNAi Library (DRSC 2.0) can be found at the <u>DRSC/TRiP Functional Genomics Resources</u>. The recommended by DRSC protocol was followed (77) on 7 series of duplicated 384 well plates screened in a range of 3 weeks. Briefly, 250 ng/well (5 μl at 50 ng/μl) of each dsRNAs were distributed in 384 well culture plates in 62 plates. One plate is organized in 16 lines (A-P) and 24 columns (1-24). Each well thus possess a unique identity number consisting in the plate number followed by the well coordinates. The dsRNA has a unique identification number (DRSC#####). A *WellMate* dispenser (Matrix Technologies) automate distributes the *automiG* S2R+ cells culture suspension in the 384 well-plates (25.000 cells at a concentration of 2,5 10^6^ cells/ml). After an hour dsRNA internalization into the cells, 30 μl of 10% serum was added. The cells were incubated for 4 days with the dsRNA before the expression of *automiG* in order to allow a complete internalization of the dsRNAs, the degradation of the target mRNAs and the catabolism of the corresponding protein. At day 5, the *automiG* expression was induced with 600 μM of CuSO_4/_well (5 μl at 6mM). After 24 hours of *automiG* induction, the cells were imaged on an *Opera* confocal microscope (Evotec Technologies, Perkin Elmer) with automated acquisition system allowing fast imaging of the epifluorescence of the cells in each well of a big number of plates. In addition, Analyst®GT multimode reader (Molecular Devices) – plate reader allowing the fast and sensible read of plates per group of 40 plates was used. A filter controls the excitation and emission wavelengths for fluorescence detection.

A validation screen was performed in triplicate using the same conditions as those used in the primary screen described above. After a 48 hours incubation period, the plates were centrifuged for 1 minute at 800 g and the culture medium was carefully removed. 25 µl of cracking buffer (125 mM Tris pH6,8, 5% ß-mercapto-ethanol, 2% SDS, 4 M urea) was added in each well and 8 µl of protein extracts were analyzed by western-blot using anti-GFP antibody as shown in Figure. 2A. Further information is available upon request and at the *<u>DRSC/TRiP Functional</u> <u>Genomics Resources</u>*, Harvard University. Database for storage and analysis of the results obtained with each RNAi screen performed at the DRSC gives access to the correspondence of the wells of the screen plates and the DRSC identity of the dsRNAs contained in these wells. It equally contains information for the dsRNAs as their quality control or the prediction of their specificity for the targeted gene.

### Amino Acid conservation and phylogenetic analysis

Sequence alignments and visualization were done in Kalign (www.ebi.ac.uk/Tools/msa/kalign/) and Unipro UGENE 1.32.0. Percentage of amino acid (aa) identities and coverages between CG7009, CG5220, TRM7, and FTSJ1 proteins were determined on the Ensembl project website (www.ensembl.org). For phylogenetic analysis, protein alignments were performed using mafft v7.407 with default parameters (78). Removal of positions with more than 50% of gaps was obtained by using trimal v1.4 (79). Phylogenetic analysis was performed using raxml v8.2.12 (80) under the PROTGAMMALG model by combining a rapid bootstrap analysis (100 replicates) and search for the best ML tree (−f a option).

### Total RNA Extraction for MALDI-TOF and RiboMethSeq analysis

3-5 days old females and males were homogenized on Precellys 24® tissue homogenizer (Bertin Technology) in 1 mL TRI-reagent (Sigma Aldrich). Total RNA from 20 mL of the fly lysates, was extracted with 8 mL of Chloroform and precipitated with 2/3 volumes of Isopropanol. The pellets were air dried and resuspended in RNase-free water.

### Purification of tRNA^Phe(GAA)^

Total RNA preparations (∼7 mg) were supplemented with LiCl to a final concentration of 0.8 M and incubated overnight at 4°C to precipitate high-molecular mass molecules. The precipitate was eliminated by centrifugation and the supernatant was supplemented with two volumes of 100 % ethanol and incubated at −20°C for two hours to precipitate small RNAs. After centrifugation, pelleted small RNAs were washed twice in 70% ethanol and resuspended in one ml of RNase-free water. tRNAs were further purified using the NucleoBond RNA/DNA 400 kit (Macherey-Nagel) following manufacturer’s instructions, except that the elution step was performed with 5 ml of 100 mM Tris-acetate (pH 6.3); 15 % ethanol and 600 mM KCl. Eluted tRNA were ethanol precipitated and resuspended in one ml of RNase-free water. Purification of tRNA^Phe(GAA)^ was performed using a 5’ biotinylated complementary oligonucleotide (5’-biotin-TGGTGCCGAAACCCGGGATTGAACCGGGG-3’) coupled to Streptavidin Magnesphere Paramagnetic particles (Promega). Annealing of specific tRNA was performed in 1 x TMA buffer (Tris-HCl pH 7.5 10 mM, ethylenediaminetetraacetic acid (EDTA) 0.1 mM, tetramethylammonium chloride 0.9 M) after heating the mixture at 95°C for three minutes followed by cooling to 60°C for 30 minutes. Paramagnetic particles were washed three times with 1 x TMA buffer and specific tRNA-Phe^GAA^ was recovered by heating the final suspension at 95°C for 3 minutes. tRNA-Phe^GAA^ was desalted and concentrated four times to 50 µL using Vivacon 500 devices (Sartorius; 2000 MWCO) using 100 mM ammonium acetate (pH 5.3) as a final buffer. The average yield obtained from 7 mg of total RNA was around 2 to 7 μg of purified tRNA-Phe^GAA^*. Note*: if used for RiboMethSeq, LiCl was washed away because of its interference with adaptor ligation during the cDNA library preparation.

### MALDI-TOF analysis of digested tRNA^Phe(GAA)^

For mass spectrometry analysis, about 500 ng of tRNA^Phe(GAA)^ were digested with 100 units of RNase T1 (Sigma) in a final volume of 10 µL at 37°C for 4 h. RNase T1 cleaves the phosphodiester bond between the 3’-guanylic residue and the 5’-OH residue of adjacent nucleotides and generates 3’-phosphate nucleosides. One microliter of digest was mixed with 9 µL HPA (40 mg/ml in water: acetonitrile 50:50) and 1 µL of the mixture was spotted on the MALDI plate and air-dried (“dried droplet” method). MALDI-TOF MS analyses were performed directly on the digestion products using an UltrafleXtreme spectrometer (Bruker Daltonique, France). Acquisitions were performed in positive ion mode.

### RiboMethSeq

RiboMethSeq analysis of *D.melanogaster* tRNAs was performed as described in (81). Briefly, tRNAs extracted from whole flies were fragmented in 50 mM bicarbonate buffer pH 9.2 for 15 min at 95°C. The reaction was stopped by ethanol precipitation. The pellet was washed with 80% ethanol and sizes of generated RNA fragments were assessed by capillary electrophoresis using a Small RNA chip on Bioanalyzer 2100 (Agilent, USA). RNA fragments were directly 3’-end dephosphorylated using 5 U of Antarctic Phosphatase (New England Biolabs, UK) for 30 min at 37°C. After inactivation of the phosphatase for 5 min at 70°C, RNA fragments were phosphorylated at the 5’-end using T4 PNK and 1 mM ATP for 1h at 37°C. End-repaired RNA fragments were then purified using RNeasy MinElute Cleanup kit (QIAGEN, Germany) according to the manufacturer’s recommendations. RNA fragments were converted to library using NEBNext® Small RNA Library kit (ref#E7330S, New England Biolabs, UK or equivalent from Illumina, USA) following the manufacturer’s instructions. DNA library quality was assessed using a High Sensitivity DNA chip on a Bioanalyzer 2100. Library sequencing was performed on Illumina HiSeq 1000 in a single read mode for 50 nt. Primary analysis of sequencing quality was done with RTA 2.12 software, to insure >Q30 quality score for >95% of obtained sequences.

Following SR50 sequencing run, demultiplexing was done with BclToFastq v2.4, reads not passing quality filter were removed. Raw reads after demultiplexing were trimmed with Trimmomatic v0.32 (82). Alignment to the reference tDNA sequences was performed with bowtie 2 ver2.2.4 (83) in End-to-End mode. Uniquely mapped reads were extracted from *sam file by RNA ID and converted to *.bed format using bedtools v2.25.0 (84). Positional counting of 5’-and 3’-ends of each read was performed with awk Unix command. Further treatment steps were done in R environment (v3.0.1). In brief, 5’-end and 3’-end counts were merged together by RNA position and used for calculation of ScoreMEAN (derived from MAX Score described previously), as well as Scores A and B (53) and MethScore (54). Scores were calculated for 2 neighboring nucleotides. Profiles of RNA cleavage at selected (candidate and previously known) positions were extracted and visually inspected.

### Northern blot

For Northern blot analysis of tRNA, 10 µg of total RNA from adults flies were resolved on a 15% urea-polyacrylamide gel (Biorad), transferred to Hybond-NX membrane (GE Healthcare) and EDC-crosslinked (Sigma Aldrich). The membranes were probed with 5′-^32^P end-labeled DNA oligonucleotides using T4 polynucleotide kinase (Fermentas). Hybridization was performed overnight at 37°C in PerfectHyb Plus (Sigma) hybridization buffer. Probe sequences are available in the *Primers and Probes section*. More details on NB procedure are available in (43).

### RNA Interference in S2R+ cells

Double-stranded RNAs (dsRNA) were synthesized by *in vitro* transcription (MEGAscript® T7 Kit, Ambion) of PCR products amplified from *w^1118^* genomic DNA using primers flanked by T7 promoters. Sequences of amplicon templates for dsRNA production are available from the *Drosophila* RNAi Screening Center (http://www.flyrnai.org/cgi-bin/RNAi_gene_lookup_public.pl). (*eg*: Ago2 DRSC10847, CG7009 DRSC39198). PCR products for T7 transcription of *fushi tarazu* (Ftz) and Firefly luciferase dsRNAs were amplified using primers: T7_Ftz_FW and T7_Ftz_Rev and T7_F_Luc_FW and T7_F_Luc_Rev, respectively (*Primers and Probes section* and (43)).

### S2R+ cell transfection

100 µL of cells at 10^6^ cells/ml resuspended in Schneider’s *Drosophila* medium (GIBCO-Invitrogen) were plated in 96-well plates. Cells were transfected with dsRNA or co-transfected with dsRNA and the corresponding sensor using Effectene (Qiagen) following the manufacturer’s instructions. 30 minutes after transfection 50 µL Schneider’s *Drosophila* medium (GIBCO-Invitrogen), completed with 10% heat-inactivated fetal calf serum, 100 U/ml penicillin and 100 mg/ml streptomycin were added. Cells were grown at 23°C without CO_2_. After 24 to 48 hours, CuSO_4_ was added to a final concentration of 600 µM and GFP fluorescence was followed using an inverted epifluorescence basic microscope. For Ago2-mediated miRNA pathway involvement (*automiG*), cells were co-transfected with 0.1 µg of *automiG*-vector and 0.32 µg of dsRNA targeting either Ago2, CG7009 or Ftz, Dcr1, Dcr2, Drosha, Ago1. 48 hours later the *automiG* promoter was induced by adding CuSO_4_ to a final concentration of 600 μM (more details in (43)).

For the luciferase assay experiment, S2R+ cells were treated for 4 days with dsRNA inactivating specifically the indicated genes. Cells were co-transfected with two plasmids expressing the Firefly and Renilla luciferases in addition to a dsRNA against Firefly. Luciferases activities were measured 48 hours after transfection. The averages of the activity ratios from Firefly/Renilla luciferases from three independent biological replicates were plotted normalized to the average of a control dsRNA (GFP) which was set to 1 (+/− the standard deviations). * indicates p< 0.05 in a Student’s T-test.

### Western Blot

Expression of GFP was analyzed in *automiG* induced cells by western blotting using mouse anti-GFP (Roche^®^) and anti-Mbf1 antibodies (43) as loading and transfer control. 72 hours after the dsRNA transfection and *automiG* vector induction, the culture medium was removed and 80 µL of Sample Buffer Laemmli 2X (Sigma^®^) was added in each well. The samples were boiled (95°C) and 18 µL were loaded onto a 4-20% Mini-PROTEAN^®^TGX^™^ 12 well-gel (Biorad^®^). After transfer onto a PVDF (Amersham Hybond, GE Healthcare) or nitrocellulose membrane, membranes were blocked in 5% milk, dissolved in 1X TBS-T (20 mM TRIS-Base, 150 mM NaCl, Tween-20 (Polyoxyethylene sorbitane monolaureate) to 0.05%) and incubated overnight with anti-GFP (1:2.000) or anti-Mbf1 (1:10.000) antibodies diluted in the blocking solution. After three times 15 minutes washes, appropriate secondary antibody (1:10.000) coupled to alkaline phosphatase (Promega) was added and incubated for one hour at room temperature. Detection was performed using BCIP (5-bromo-4-chloro-3-indolyl-phosphate) and NBT (nitro-blue-tetrazolium, (ThermoFischer) reagents diluted in AP buffer (100 mM Tris-HCl pH 9.5, 100 mM NaCl, 5 mM MgCl_2_).

### automiW

Experiments with the *automiW* eyes sensor were performed as described in (47). Eye images were acquired with an Axio-ApoTome (Zeiss) and ZEN2 software.

### DCV injection

Flies with the following genotypes were subjected to intra-thoracic injection with the *Drosophila C* virus (DCV): *CG7009^e02001^*/+ (controls). *CG7009^e02001^*/*Def3340* or *CG7009^e02001^*/*CG7009^e02001^* (*CG7009* mutants). *CG5220^K28A^*/*CG5220^K28A^* (*CG5220* catalytically dead mutant). 2 to 4 days old flies were divided in tubes of 10 (5 males + 5 females) and 20 flies from each genotype were injected with DCV while 20 other flies were injected with adult Injection Buffer (controls), containing 10 mM Tris pH 6.3 and 1mM MgCl_2_. Each fly was injected with 4.6 nL of DCV concentration of 2×10^6^ PFU/ml (9.2 PFU/injection). Intra-thoracic injections were made using the Drummond Automatic Nanoliter Injector “NANOJECT II”. After injection, flies were kept at 25°C. Three to four days after the injection and Before death, 3 injected flies from each genotypes and conditions (+ or − DCV) were frozen at −20°C. 3 flies from each condition were then crushed with a pestle in TRI-Reagent (Sigma Aldrich) and total RNA was extracted as described above. DNase digestion and RT-qPCR were carried out as described with DCV_FW and DCV_Rev specific primers and Rp49 for normalization (*Primers and Probes section*).

### CRISPR/Cas9-mediated genome editing and genotyping

Mutant alleles for *CG5220* were generated using CRISPR/Cas9-mediated editing in *Drosophila* as previously described (85). The *CG5220*^K>A^ allele was obtained using the gRNA (guide RNA) sequence 5’-CTTCGAGCAACTTGAAGGCACTCC-3’. ssODN: donor template K to A mutation in CG5220 was 5’-TTCATATATTTATTTACAATGGGGAAAACATCAAAGGACAAAAGAGATATCTATTACCGAC AAGCCAAAGACGAAGGCTGGAGGGCGAGGAGTGCCTTCGCGTTGCTCCACGTGGACGAAGCCTACGGAATTCTAA-3’. A mix of 150 ng/µl of the vector and ssODN was injected in pre-blastoderm *w^1118^* embryos. Screening for flies containing the substitution was carried out by PCR on F1 males derived from crossing of injected individuals with a balancer stock. The obtained sequences correspond to fragments covering 438 bp of the *CG5220* gene. Fly stock denominations are *CG5220^K>A^*. Visual sequence analysis was carried out using *4Peaks* and *ApE* softwares.

### RT-qPCR

Whole flies or dissected ovaries were crushed with a pestle in TRI-Reagent (Sigma Aldrich). After DNase digestion of total RNA using the TURBO DNA-free™ Kit (Ambion), 500 ng were used in a reverse transcription reaction with Random Primers (Promega) and SuperScript® II Reverse Transcriptase (Invitrogen). The cDNA was used to perform qPCR on a CFX96 Touch™ Real-Time PCR Detection System (Bio Rad) using target-specific primers. Rp49 was used for normalization (*Primers and Probes section*). The analysis was performed using ΔΔ Ct, error bars represent SD of three biological replicates. Statistical analysis using a Student’s T-test was performed and *p*-values were determined.

### Production and affinity purification of recombinant fusion proteins

Glutathione *S*-transferase (GST) fusion constructs were generated by PCR amplification of full-length cDNAs of CG7009 available from BDGP (#SD16956) using standard PCR with VENT polymerase (New England BioLabs). Products were cloned between the EcoRI and NotI restriction sites of the pGEX-4T1 (GE Healthcare) vector using primers CG7009_EcoRI_ATG and CG7009_NotI_Stop. Amplification of full-length cDNAs of *CG33172* (clone MIP10235 in BDGP) was done using standard PCR techniques using Q5 high fidelity DNA Polymerase (*New England BioLabs*). Amplification products were cloned between the HindIII and NotI restriction sites of the pET-28a (Novagen) vector (modified to contain FLAG peptide) using the primers CG33172_Hind_ATG and CG33172_NotI_Stop. Competent bacteria TOP10 (Invitrogen) were transformed by heat-shock with 100-200 ng of plasmid DNA according to each manufacturer’s instructions. After expression on the corresponding antibiotic resistance genes by incubation for 0.5-1,5 hours at 37°C under agitation of 250 rpm, 1/10 and 9/10 of the transformed bacteria were plated on LB agar plates, supplemented with the corresponding antibiotics. GST fusion proteins were expressed in Escherichia coli BL21 Star (DE3) (Invitrogen) or C41 (86) and purified over glutathione-coupled resin (Pharmacia) as previously described (87, 88). The same protocol was used for purification of pET-28a Flag fusion proteins. Bound peptides were eluted with 400 μg/ml Flag peptide (Sigma) in BC100 buffer for 20 min on ice.

### In vitro interaction of GST-CG7009 and FLAG-CG33172

Briefly, GST-alone (control) or fusion proteins GST::G7009 (pGEX4T1-CG7009) and FLAG::CG33172 (pET28a-FLAG-CG33172) were co-expressed in C41 (86) bacteria and purified over Flag-coupled resin (Sigma). Bound proteins were washed 3 times in 500 mM KCl and eluted on Bio-spin disposable chromatography columns (Bio-Rad) with flag peptide as described in (88). Western blot of the immunoprecipitated recombinant proteins was performed as described in the above section using anti-GST HRP (horseradish peroxidase) conjugate (1:10,000 Amersham GE Healthcare) for 60 minutes at room temperature under agitation. HRP was detected by enhanced chemiluminescent (ECL).

### RNA-seq on S2R+ cells

Knock downs for *CG7009* and *LacZ* (control KD) in S2R+ were performed in a serum-free medium using 7.5 μg of dsRNA per 10^6^ cells and stopped 2 hours after cell starvation with the addition of the serum-supplemented medium. dsRNA treatment was repeated after 48 hours. Cells were collected and total RNAs were extracted 96 hours after the first treatment. Libraries were prepared using the Illumina TruSeq Sequencing Kit by following the manufacturer’s protocol for paired-end read mode and directional sequencing on an Illumina NextSeq 500 with a read length of 42 bp.

### Computational analysis of RNAseq experiments

Basecalling and demultiplexing were performed using bcl2fastq (v2.19). Individual samples were mapped using STAR (89), v2.5.2b) against ensembl release 90 of the Drosophila melanogaster genome (BDGP6). Gene counts were derived using featureCounts ((90), v. 1.5.1). RNA-Seq Analysis Differential expression analysis was performed using Bioconductor v2.38/ DESeq2 v1.18.1 (91, 92). Genes were called sig. diff. expressed with an FDR below 5%. The sample Ctrl_3 was excluded as an outlier from the differential expression analysis.

### Imaging

Ovaries and eyes images were acquired with a WILD M3Z (Leica) binocular combined with a Q IMAGING Color 12 bit (Q27959) camera and QCapture Pro software.

### Weighing

Average weight for flies in milligrams (mg) was calculated for 3-days-old flies measured on precision balance (>= 0,001g) after dehydration of the bodies. n >=14, where each n is the mean value of 10 flies measured in a batch of 10. *p*-value<0,001 in a Student’s T-test.

### Lifespan assays

2-3-day-old male flies Da-GS; UAS-RNAi *CG5220*, *CG7009* were kept at 25 °C in vials with standard medium complemented (RU200) or not (RU0) with RU486. RNAi transgene induction using the Da-Gal4-GS (GeneSwitch) lines was described in (93). Briefly, the GeneSwitch protein (GS) is a GAL4 modified protein that recognizes and activates UAS-dependent transgenes only in the presence of RU486 added into *Drosophila* medium food. The number of flies tested was 3 times 10 flies. To monitor survival rates over time, flies were counted and transferred into a new tube every 2-3 days. Constitutive expression of *CG5220*, *CG7009* KD transgenes was induced by RU486 exposure (20 mg/ml) during adulthood.

### Drosophila stocks

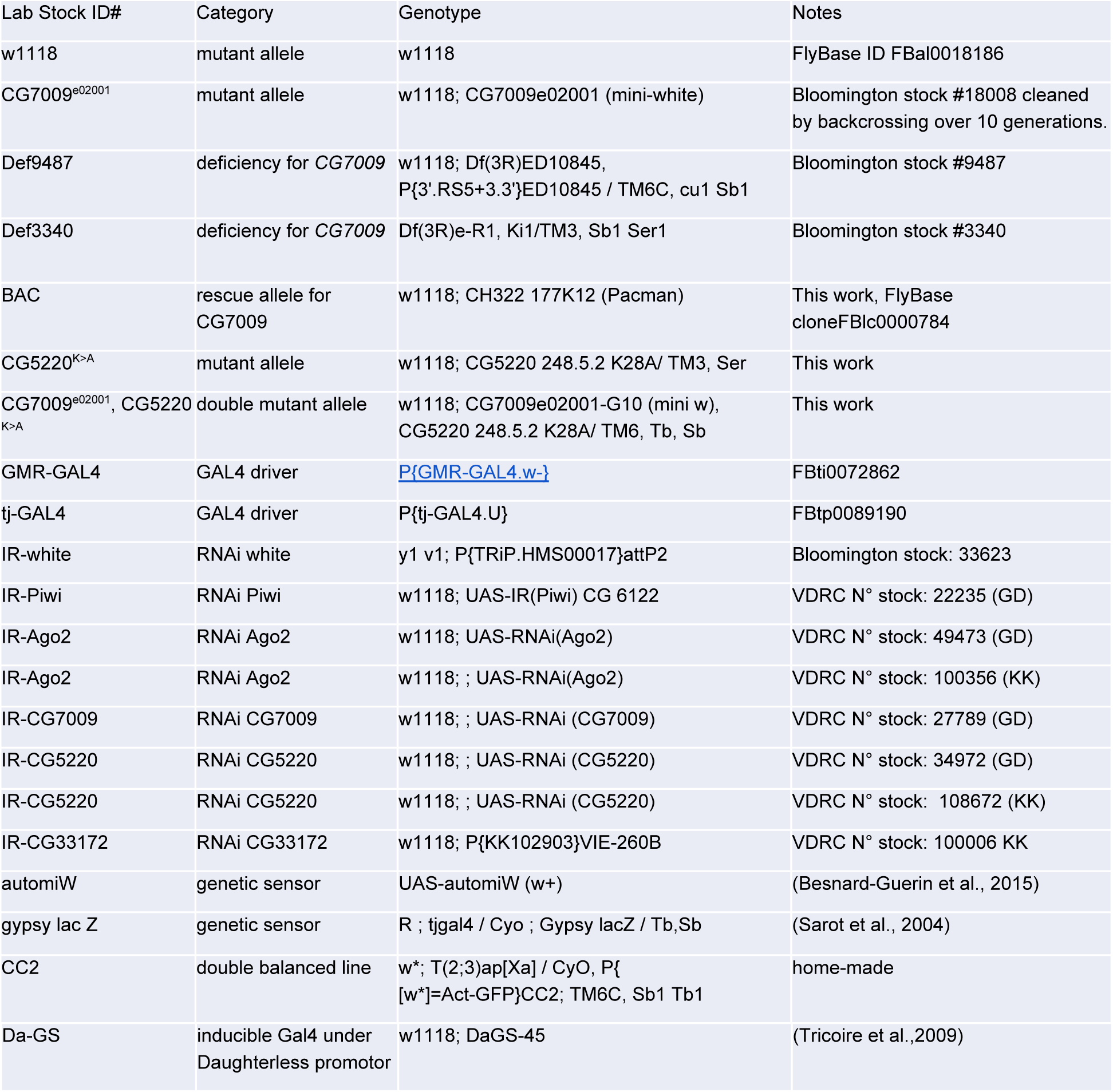

### Primers and Probes

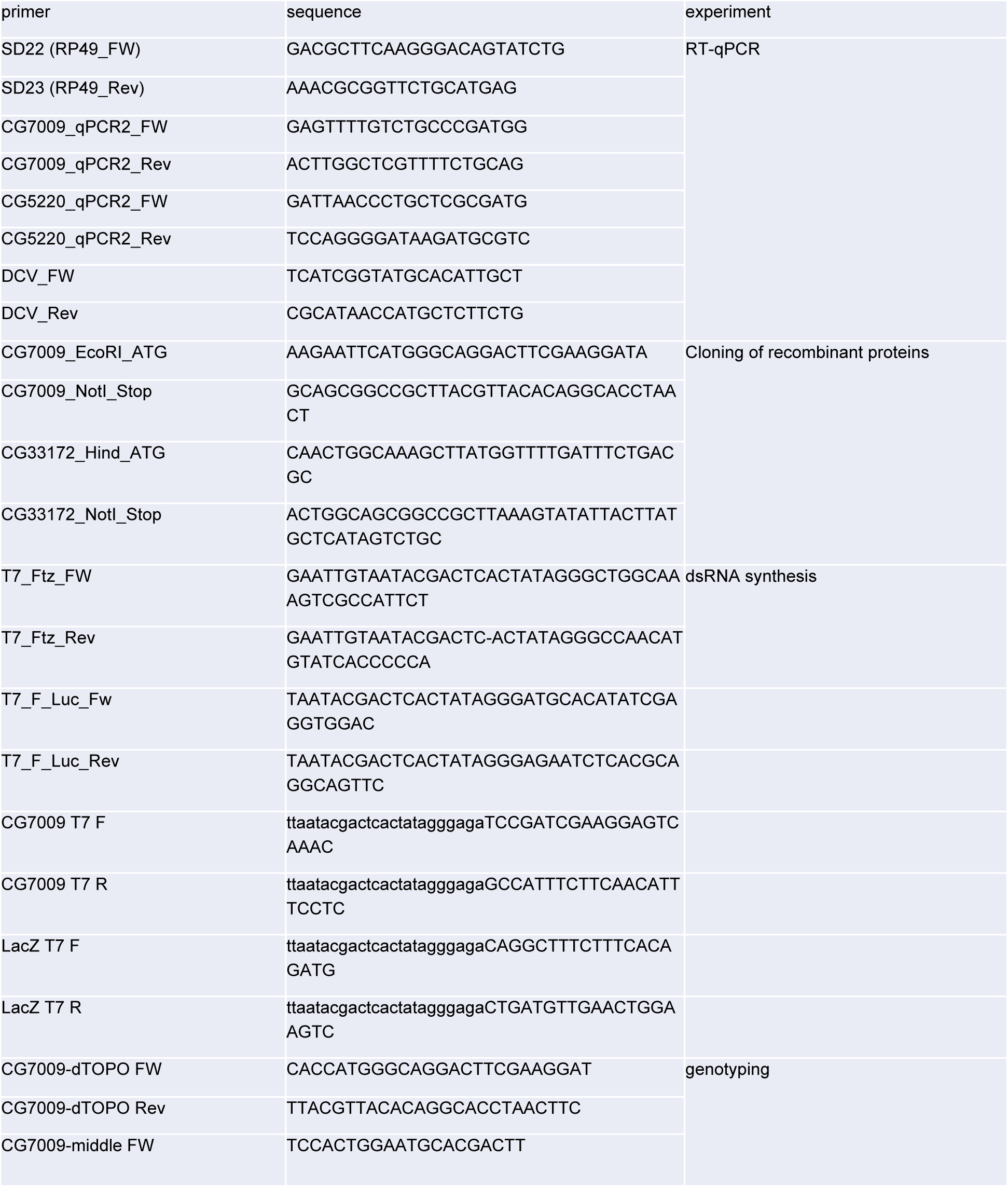
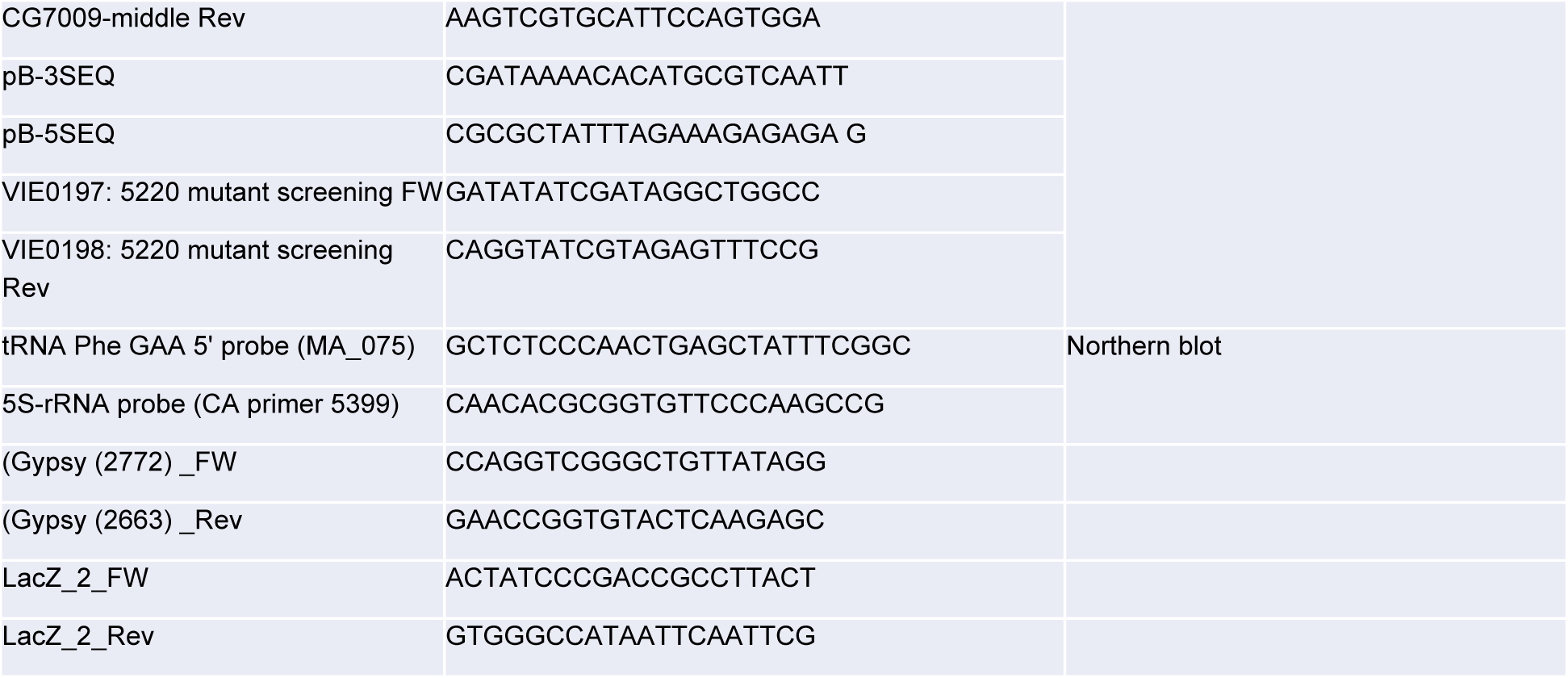

## Data availability

The RNA sequencing data discussed in this publication have been deposited in NCBI’s Gene Expression Omnibus (94) and are accessible through GEO Series accession number GSE134354 (https://www.ncbi.nlm.nih.gov/geo/query/acc.cgi?acc=GSE134354).

## Supporting information

RNAseq analysis

## Acknowledgements

We would like to thank the DRSC/TRiP Functional Genomics Resources for the DRSC 2.0 genome-wide screening library. Bloomington *Drosophila* Stock Center for fly reagents; the *Drosophila* Genomics Resource Center at Indiana University for plasmids; Jean-Luc Imler for sharing DCV preparations; Romain Derelle for the phylogenetics analysis; Valérie Biou and Bruno Miroux for the C41 bacteria; members of the TErBio laboratory and ARTbio bioinformatic platform as well as Stephan Eberhard for helpful discussions and careful reading of the manuscript; Josette Pidoux, Elma Shalatzi, Marius van den Beek, Souraya Kouider, Elie Makardijian and Ludivine Roumbo for experimental help. Support by the IMB Genomics Core Facility and the use of its NextSeq500 (INST 247/870-1 FUGG) is gratefully acknowledged. We also thank Nastasja Kreim from the Core facility Bioinformatics for her great support. C.C. received financial support from CNRS and Sorbonne Université, Fondation Maladies Rares (FMR), IBPS grant “*Action Initiative 2018*”, the COST action EPITRAN CA16120. M.T.A. and D.G.D. had PhD fellowships from the Ministère de la Recherche et de l’Enseignement Supérieur at the doctoral school Complexité du Vivant (ED515). We also thank the Fondation ARC pour la Recherche sur le Cancer for funding support to M.T.A. and D.G.D. (4th year PhD), the “Réseau André Picard”, the “Société Française de Génétique” for traveling fellowship grants to M.T.A. and D.G.D., the COST action “EPITRAN” CA16120 for traveling fellowship grants to D.G.D. and C.C. J-Y.R. Y.M. M.S. and C.C. are members of the European EPITRAN COST Action (CA16120).

## Author contributions

M.T.A, D.G.D., B.d.S., S.S. and C.C. performed all experiments in regard to fly genetics, life span measurements, *CG7009* allele characterizations and RNAs and recombinant proteins preparation. M.S designed the strategy and created the *CG5220^K>A^* allele. C. Achour generated and screened the *CG7009, CG5220* double mutants. C.J., C.A. and C.C. designed and performed the genetic and secondary sensor screens. C.G. and D.B. purified tRNAs. V.G. performed Mass Spectrometry and analysed the data with D.B. and C.G. V.M. and V.B.I. performed RiboMethSeq. RiboMethSeq computational analyses were performed by Y.M. J.Y.R. and T.L. performed transcriptome sequencing libraries preparation and computational analysis was performed at the IMB Genomics Core Facility and ARTbio with the help of C.A. M.S. performed northern blot experiments. C.Achour, M.T.A. and C.C. performed GST pull down experiments. M.T.A., D.G.D., Y.M. and C.C. prepared Figures and Tables. C.C. conceived the project. M.T.A., D.G.D., L.T., J.Y.R., Y.M., M.S. and C.C wrote the manuscript.

## Supplementary Figure Legend

**Figure S1.**
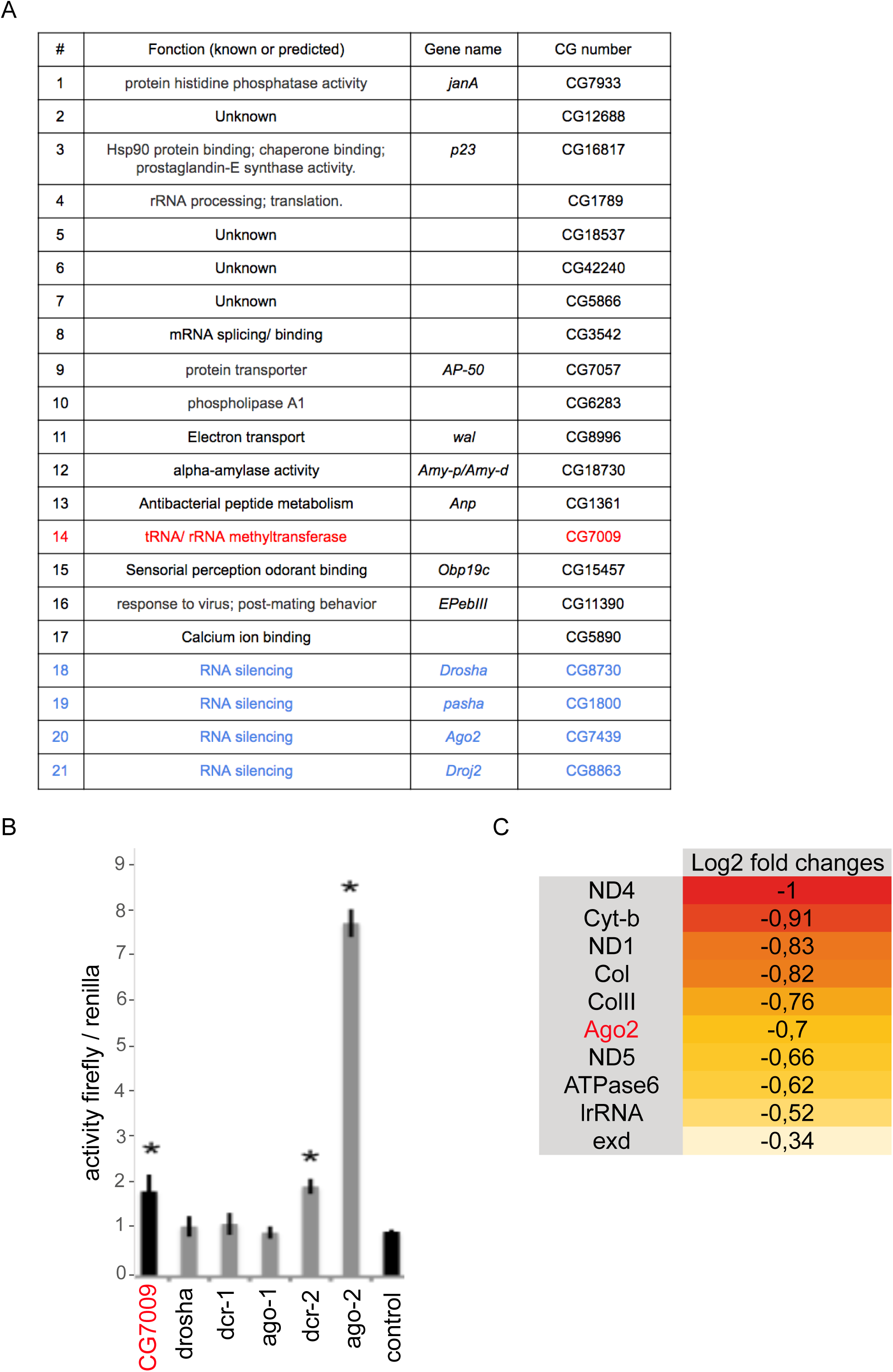
**A**, List of the validated genes in the RNAi genome-wide screen. Indicated are the genes that scored positively in the screen, their described or predicted function in FlyBase, the gene name (when available), and the CG number, according to FlyBase. Blue colored names indicate genes already known in RNA silencing pathways. Red point to CG7009. **B**, siRNA sensor (44, 45) reveal failure in Dcr2/ Ago2 dependant siRNA pathway. After 4 days of treatment with a dsRNA specifically inactivating the indicated gene, S2 cells were co-transfected with 2 vectors expressing the luciferases firefly and Renilla (control) in addition to dsRNA firefly (dsRNA Fluc). Firefly and renilla activities were measured 48 hours after transfection. The averages of the ratios of firefly/renilla activity for 3 independent experiments ± s.d are represented, the average for the dsRNA GFP (control) being set at 1. * indicates p<0.05 in a Student’s T-test. **C**, Ago2 mRNA is downregulated upon *CG7009* KD. The heat map shows the top 10 differentially expressed genes in S2 cells upon *CG7009* KD versus control (*LacZ* KD) in a RNA sequencing experiment. log2 Fold Change is indicated (padj = FDR-adjusted *p*-value for Ago2 is 7.73e^−118^). Red to yellow heat map colored table (Red: most down-regulated; yellow, less down-regulated). KD of *CG7009* led to the deregulation of 173 genes including 45 up and 128 down. 110 genes were deregulated with a corrected *p*value (Benjamini and Hochberg, FDR) < 0.01 (not shown).

**Figure S2.**
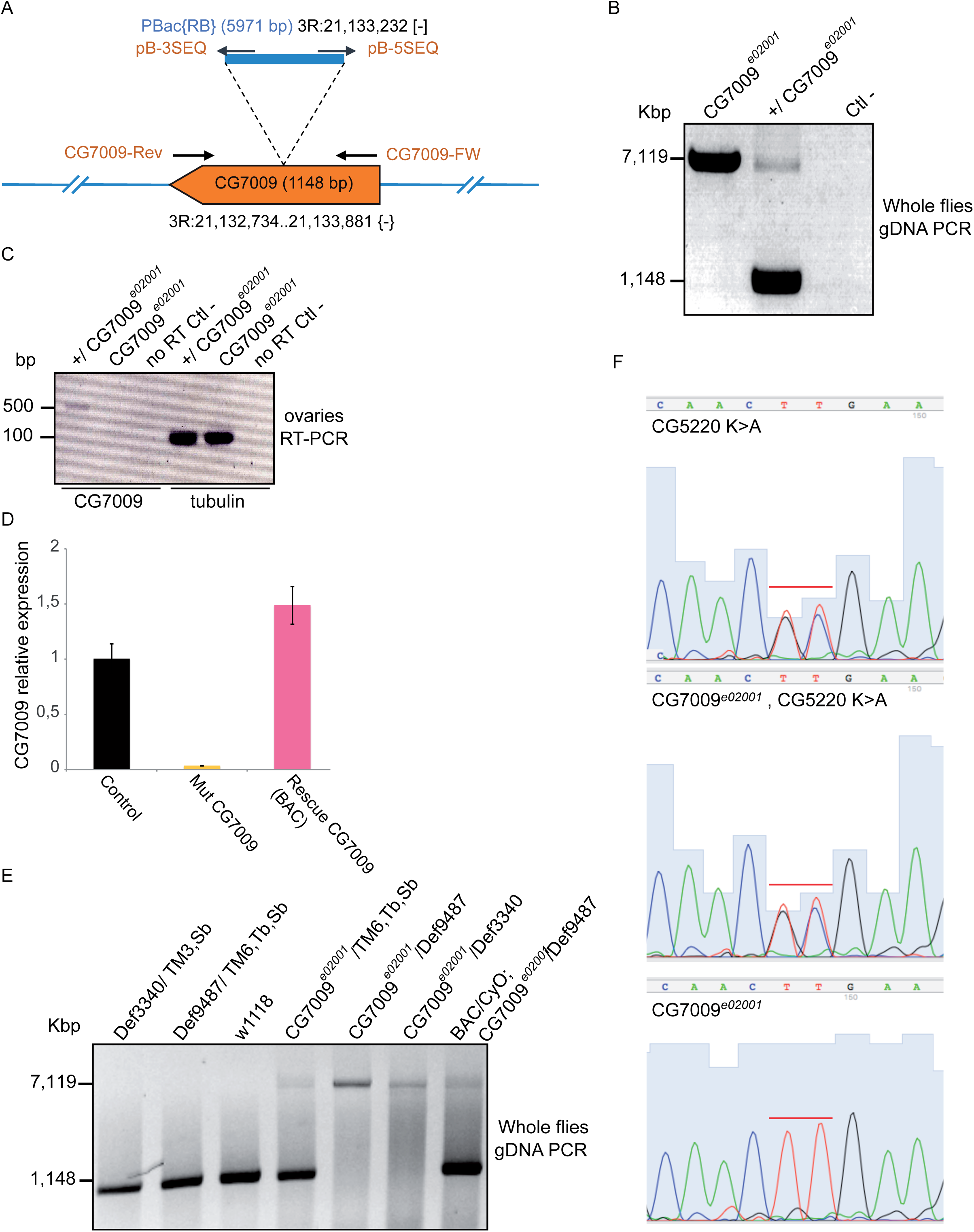
**A**, schematic representation of the *CG7009^e02001^* allele showing the genomic location of the insertion and the size of the PBac{RB} transgenic transposon. The four hybridizing oligonucleotides used in the genomic PCR analysis are represented with arrows representing their annealing locations and orientation (sequences in *Primers and Probes section*). **B**, Agarose gel electrophoretic separation of the PCR reaction made on gDNA (genomic DNA) of adult flies *CG7009* homozygous or heterozygous for the mutant allele *CG7009^e02001^* or Ctl - (no DNA) using CG7009-FW and CG7009-Rev primers. Expected sizes in base paired (Kbp) are indicated on the left. **C**, Reverse Transcription PCR (RT-PCR) on total RNA extracts from ovaries. Electrophoretic separation of the RT-PCR reaction made on total RNA of flies *CG7009* homozygous or *CG7009* heterozygous for the mutant allele *CG7009^e02001^* or no RT control (no reverse transcriptase in the RT reaction). The used primers for the PCR reaction are CG7009-middle Rev and CG7009-FW (expected product size 500nt) and tubulin primers (expected product size 150nt). **D**, Reverse Transcription qPCR (RT-qPCR) on total RNA extracts from adult females of the indicated genotypes. *CG7009^e02001^* heterozygous (Control), *CG7009^e02001^* homozygous mutant (Mut CG7009) and rescue CG7009 (BAC). **E**, Genotyping by PCR on genomic DNA of heterozygous *Def9487* and *Def3340* and *CG7009^e02001^*. BAC (rescue) / CyO; *CG7009^e02001^* / *Def9487* lines, w1118 and *CG7009^e02001^* homozygous lines. PCR on gDNA extracted from adult single flies with the indicated genotypes using primers CG7009-FW and CG7009-Rev. The band at 1148 bp corresponds to the WT *CG7009* locus, the band at 7119 bp corresponds to the mutant allele *CG7009^e02001^*, containing the inserted *PiggyBac* transposon (Supplementary Figure S2A, B). BAC: Bacterial Artificial Chromosome containing the wild type *CG7009* genomic region; CyO; TM3,Sb; TM6,Tb,Sb: balancer chromosomes; Kbp: Kilo base pairs; gDNA genomic DNA. **F**, Validation by sanger sequencing of the CRISPR/Cas9 mutants *CG5220^K>A^* and double mutant *CG5220^K>A^, CG7009^e02001^* recombination. Briefly, *CG5220* PCR fragments were amplified by PCR from flies gDNA bearing the *CG7009^e02001^* allele (giving the [w+] phenotype) and the mutations *CG5220^K>A^*. The corresponding simple mutant *CG5220^K>A^* (heterozygous for *CG5220^K>A^*) was used as positive controls and flies characterized with no *CG5220* mutation were used as negative controls (*CG7009^e02001^*). All sequencing experiments were performed on heteroallelic combinations over balanced chromosomes, explaining the double picks at the edited region. The results were obtained using 4Peaks. They correspond to a PCR products obtained using VIE0197/VIE0198 primers and sequenced with the primer VIE0198. The targeted nucleotides are indicated under the red lines.

**Figure S3.**
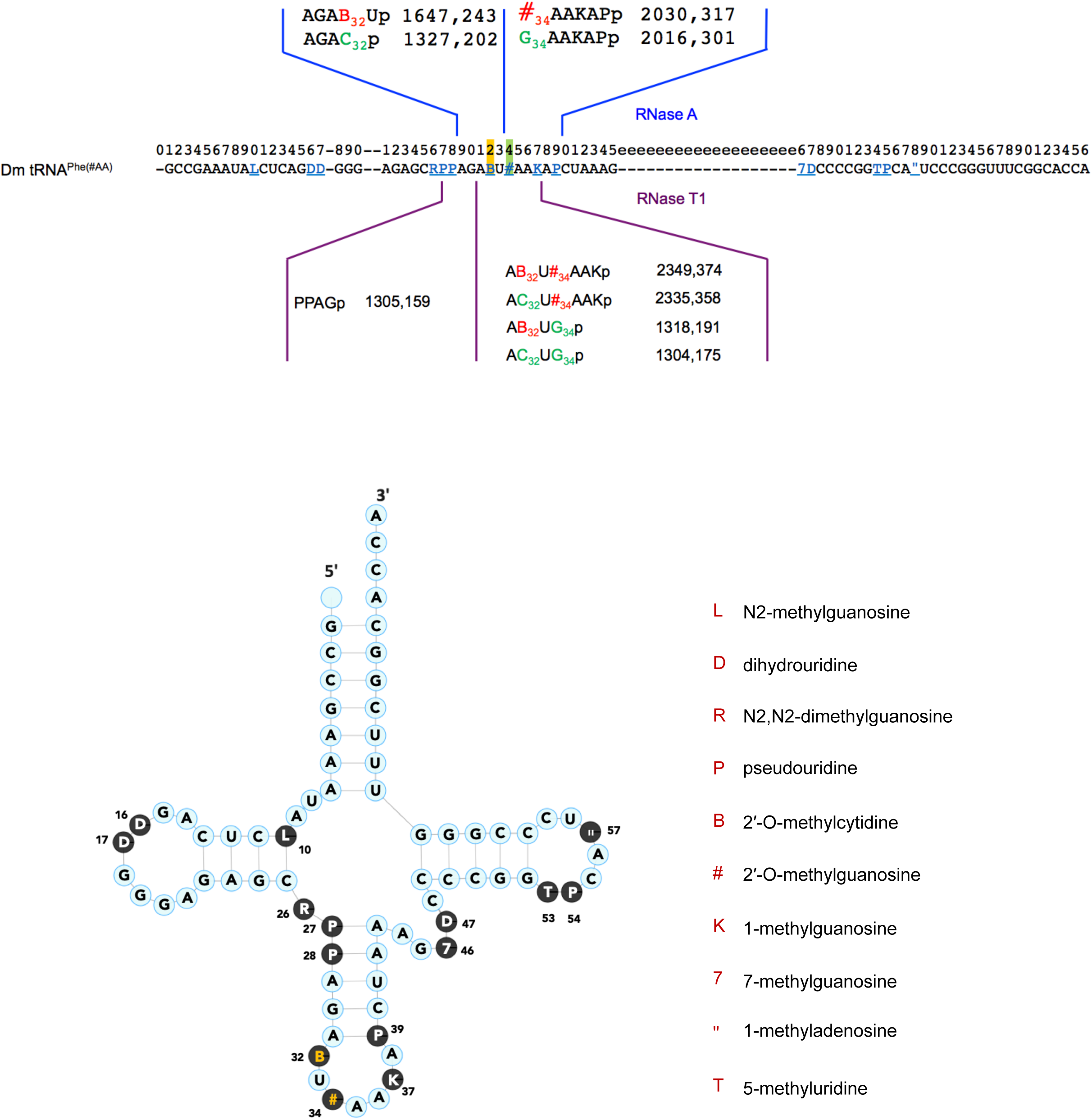
**Top:** Sequence of *Drosophila melanogaster* tRNA^Phe(GAA)^ with m/z values of fragments containing 2’-O-methyl-guanylate (#) and/or 2’-O-methyl-cytidylate in daltons. The expected fragments resulting from RNase A (top) and RNase T1 (bottom) digestion of tRNA^Phe(GAA)^ with the indicated Nm modification are zoomed in. **Bottom Left**: Schematic representation of tRNA^Phe(GAA)^ of *Drosophila*. Cm_32_ (B) and Gm_34_ (#) are indicated in yellow. **Bottom Right**: Names of the different modifications present on *Drosophila* tRNA^Phe(GAA)^ based on the <u>Modomics</u> and <u>tRNAdb</u> nomenclature.

**Figure S4.**
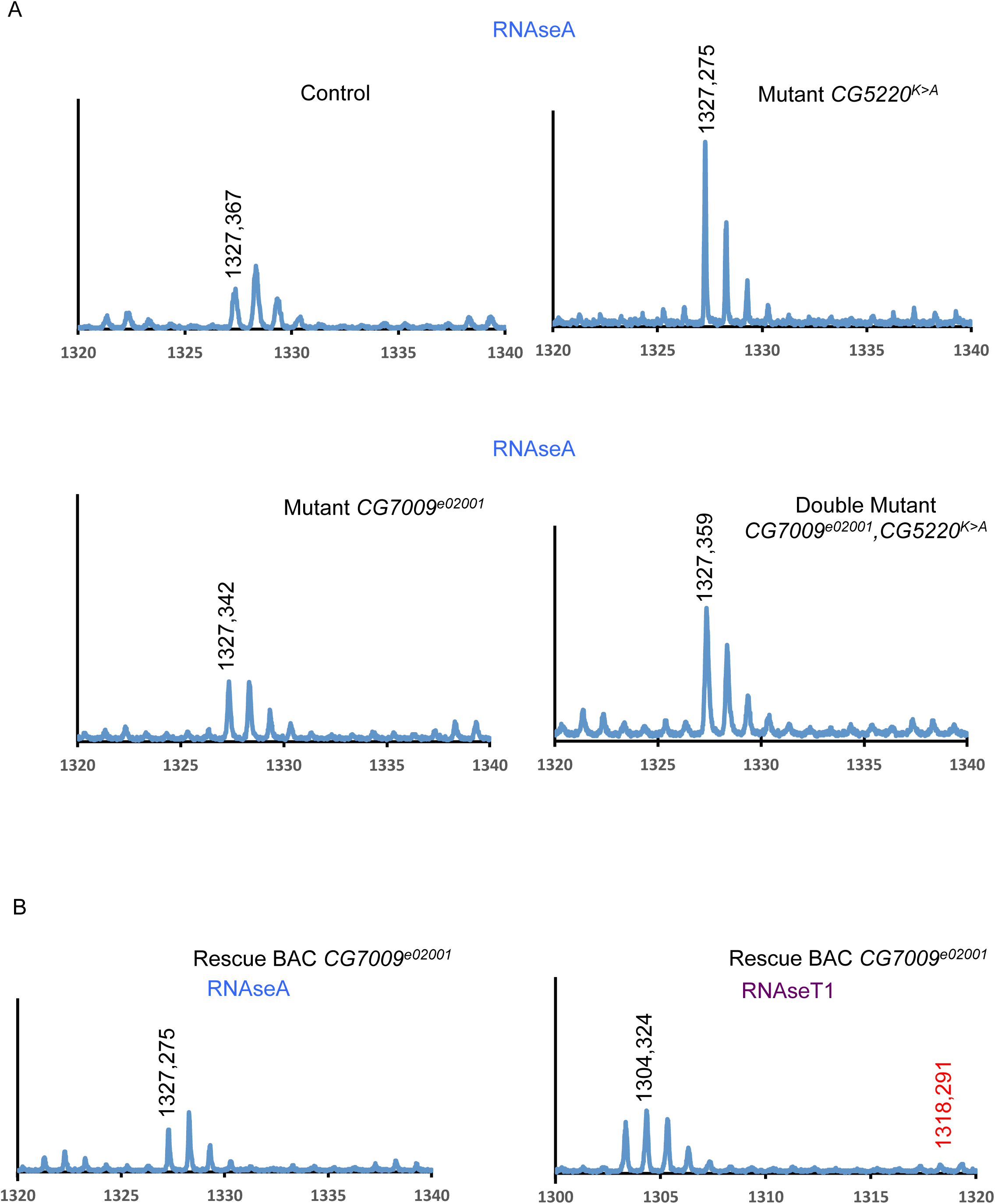
**A**, MALDI TOF-MS spectrum of fragments resulting from RNase A digestion of tRNA^Phe(GAA)^ originating from indicated genotypes. **B, Top**, MALDI TOF-MS spectrum of fragments resulting from RNase A digestion of tRNA^Phe(GAA)^ originating from indicated genotypes (homozygous adult *CG7009^e02001^* mutants rescued with one CG7009 WT copy (BAC)). **Bottom**: MALDI TOF-MS spectrum of fragments resulting from RNase T1 digestion of tRNA^Phe(GAA)^ originating from indicated genotypes (homozygous adult *CG7009^e02001^* mutants rescued with one CG7009 WT copy (BAC)). Relevant peaks are identified by their m/z values.

**Figure S5.**
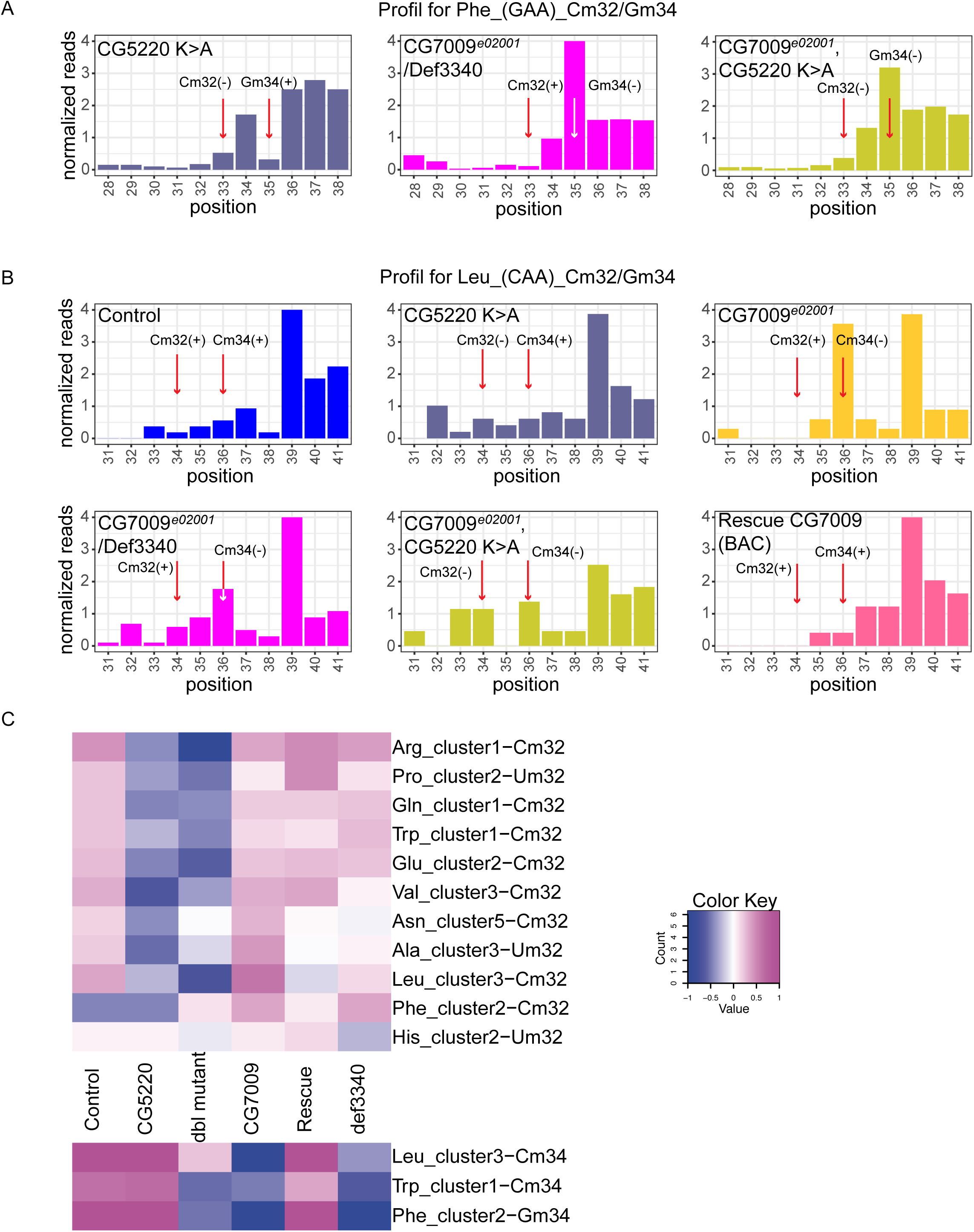
**A**, related to Figure 3C. RiboMethSeq analysis of tRNA^Phe(GAA)^ modification at positions Cm and Gm_34_. Alkaline fragmentation-based RiboMethSeq protocol was performed on total RNAs extracted from whole flies homozygous mutant for *CG5220^K>A^*, homozygous for *CG7009^e02001^*/ *Def3340* and homozygous double mutant *CG5220^K>A^*, *CG7009^e02001^* as indicated. Normalized cleavage efficiency, calculated from cumulated 5’-end and 3’-end coverage, is shown for the ± 5 neighboring nucleotides. The positions of interest (Cm and Gm) in tRNA^Phe(GAA)^ are shown by red arrows. Protection against cleavage is indicated (+) protected, (−) not protected. Protection at Cm_32_ in control flies is only moderate, indicating incomplete tRNA methylation (+low). Higher methylation levels at this positions are observed in *CG7009* homozygous and Rescue CG7009 (BAC) in Figure 3C **B**, RiboMethSeq was performed as described in Supplementary Figure S5A above on tRNA^Leu(CAA)^ on the 6 indicated genotypes. Normalized cleavage efficiency, calculated from cumulated 5’-end and 3’-end coverage, is shown for the ± 5 neighboring nucleotides. The positions of interest (Cm and Cm) in tRNA^Leu(CAA)^ are shown by red arrows. Protection against cleavage is indicated (+) protected, (−) not protected. Protection at Cm_32_ in control flies is only moderate, suggesting incomplete tRNA methylation. Protection levels clearly demonstrate that Cm_34_ is missing in *CG7009* homozygous mutant and rescued in rescue CG7009 condition while Cm is CG5220 dependent. Increased number in D-loop tRNA^Leu(CAA)^ explain the discrepancy between real ACL position (32 and 34) and sequence positions. **C**, Heatmap of normalized MethScores (ScoreC) for tRNA positions Cm/Um_32_ and Cm/Gm_34_ in different indicated fly mutant backgrounds. Pink color corresponds to overmethylation compared to the average, blue or white color to undermethylation. Simplified genotypes is indicated in the middle, affected tRNA species are given on the right. Only positions 32 and 34 affected in mutant flies are shown. Scale of the color key (−1 to +1) is indicated at the top.

**Figure S6.**
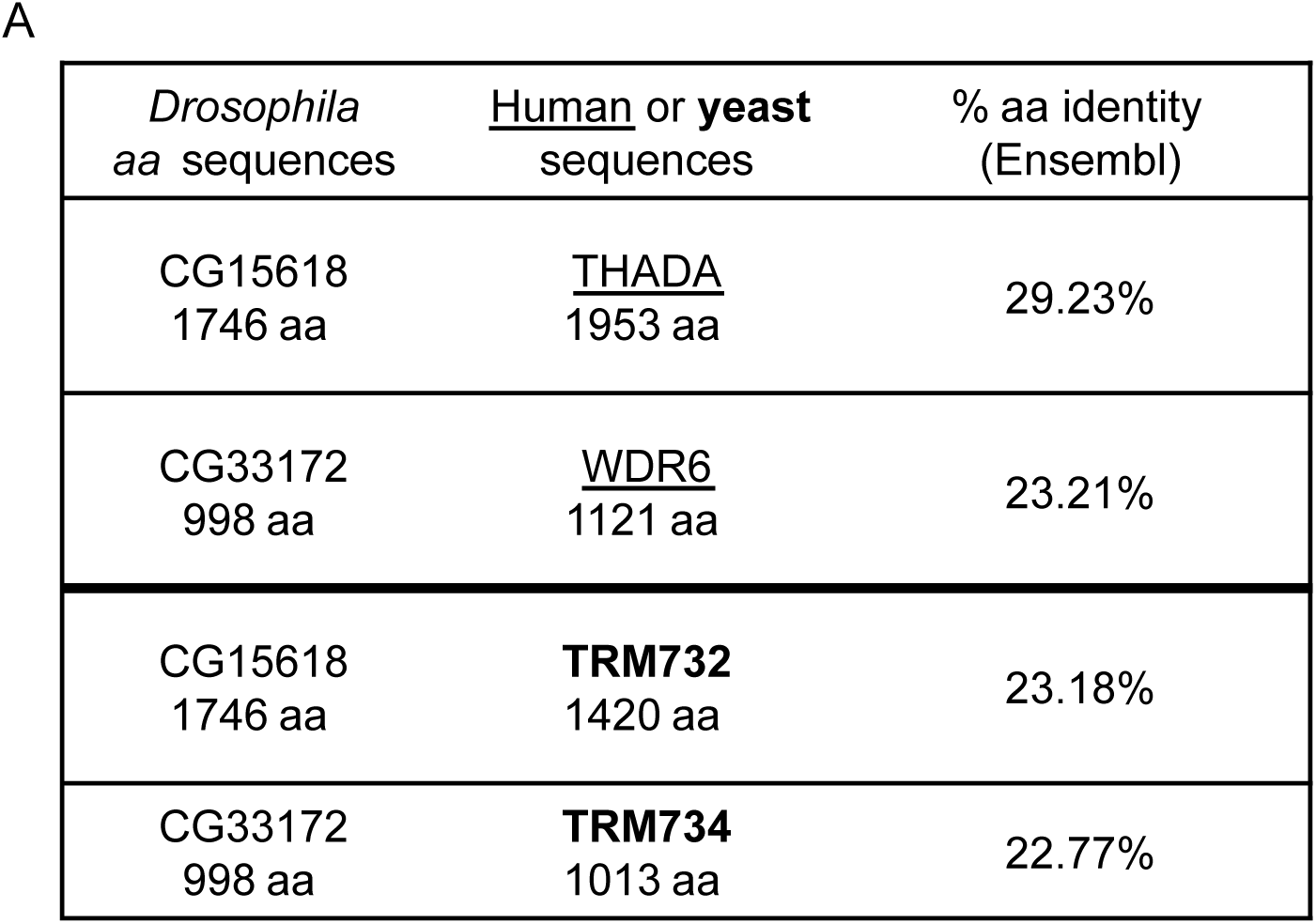
**A**, Pourcentage of amino acid (aa) identity between CG15618, human THADA and yeast TRM732, and between CG33172, human WDR6 and yeast TRM734 (RTT10). Alignment was performed using BLAST/BLAT tool at www.ensembl.org.

